# Anxious individuals are more sensitive to changes in outcome variability and value differences in dynamic environments

**DOI:** 10.1101/2024.08.25.609575

**Authors:** Brónagh McCoy, Rebecca P. Lawson

## Abstract

Anxiety is known to alter learning in uncertain environments. Standard experimental paradigms and computational models addressing these differences have mainly assessed the impact of volatility, and anxious individuals have been shown to have a reduced learning rate when moving from a stable to volatile environment. Previous research has not, however, independently assessed the impact of both changes in volatility, i.e., reversals in reward contingency, and changes in outcome variability (noise) in the same individuals. Here we use a simple probabilistic reversal learning paradigm to independently manipulate the level of volatility and noise at the experimental level in a fully orthogonal design. We replicate general increases, irrespective of anxiety levels, in both positive and negative learning rates when moving from low to high volatility, but only in the context of low noise. When low volatility is combined with high noise, more anxious individuals display negative learning rates similar to high volatility with high noise, whereas those lower in anxiety show the usual negative learning rate increase from low to high volatility. Within-individual increases in lose-shift responses from low to high noise conditions scale with levels of anxious traits, but this occurs under low volatility only. We furthermore find that people with higher anxious traits are more accurate overall and utilize a more exploitative decision-making strategy in this dynamic environment. Our findings suggest that changes in both sources of uncertainty, volatility and noise, should be carefully considered when assessing learning, particularly in relation to anxiety and other neuropsychiatric conditions, and implicate anxiety-related differences in dopaminergic and noradrenergic neurotransmitter signalling when learning in highly changeable environments.

## Introduction

Learning in uncertain environments requires appropriate and often significant adaptation. In recent years, researchers have focused on how healthy individuals and those with neuropsychiatric conditions respond to environmental volatility, by probing how individuals adjust their learning based on how changeable the environment is, i.e., how the association between a stimulus and/or action and its subsequent outcome changes over time (Behrens et al., 2007). People with anxiety disorders often experience symptoms such as a greater intolerance of uncertainty (Bishop 2007; Carleton, 2016; Goris et al., 2020; Sandhu et al. 2023), a greater sensitivity to negative events (Grupe & Nitschke, 2013), and elevated avoidance tendencies (Maner & Schmidt, 2006; Berman et al. 2010) compared to healthy controls. All of these features of anxiety have been linked to difficulties in learning under uncertainty, but the mechanisms by which uncertainty disrupts adaptive learning in anxiety is not fully understood.

To reduce uncertainty, the brain is believed to operate in a Bayesian way, evaluating the statistical structure of the environment, and continuously trying to make sense of the causes of sensory inputs (de Lange et al., 2018; Doya et al., 2007; Rao and Ballard, 1999). According to reinforcement learning (RL) theory, surprising events lead to prediction errors, which are then used to update beliefs about the structure of the environment (Rescorla & Wagner, 1972; Sutton & Barto, 1998). Learning rates indicate the extent to which prediction errors are incorporated into these updated values. Whether to ascribe surprising outcomes to expected uncertainty – inherent uncertainty about the probabilistic nature of events (i.e., noise) - or to unexpected uncertainty – a change or context switch in the stimulus we’re learning about (i.e., volatility) - is a trade-off we encounter every day (Pulcu & Browning, 2019; Bland & Schaefer, 2012; de Berker et al., 2016; O’Reilly, 2013; Yu & Dayan, 2005). Previous literature has made predictions about how learning rates may change under different types of uncertainty (Yu & Dayan, 2005). If surprising outcomes are caused by noise, i.e., substantial variability in outcomes, then current actions should be based on the average over the outcomes of many previous actions. This is equivalent to a *lower learning rate for noisier environments*, which puts less weighting on individual outcomes. If surprising outcomes are instead caused by a volatile environment, then greater emphasis should be placed on the most recent events to guide optimal choices, i.e., by *increasing learning rate*. A high learning rate may result in over-learning, with more win-stay (sticking with the same choice after receiving a positive outcome) and lose-shift (switching to a different option after receiving a negative outcome) behaviour, whereas a low learning rate may lead to poor updating and result in slow adaptation to a change in the environment (Huang et al., 2017). Many previous studies have reported an increase in learning rate for volatile compared to stable environments (Behrens et al., 2007; Browning et al., 2015; Lawson et al. 2017; Manning et al., 2017). Sensitivity to positive and negative outcomes, e.g., positive and negative learning rates, has previously been linked to win-stay and lose-shift behaviour, respectively (St. Onge et al., 2011), although other researchers suggests that RL parameters and win-stay/lose-shift behaviour may represent separate, potentially complimentary behavioural strategies (Worthy and Maddox, 2012, 2014).

Learning under uncertainty also depends on the extent to which learned values of stimuli are used to guide decision-making. This is captured by the value sensitivity (inverse temperature) parameter in RL models, capturing the difference between learned values of options, and is representative of an exploration-exploitation trade-off. Studies have shown that from childhood to adulthood, people become less exploratory in their value-based decision-making, i.e., they are more inclined to exploit learned value differences when making decisions (Nussenbaum & Hartley, 2019). Better task performance during probabilistic RL tasks has been linked to a more exploitative strategy, indicated by a higher value sensitivity parameter (Jahfari et al., 2019; McCoy et al., 2019). Increased value sensitivity has also been associated with increased pupil size during a RL task (van Slooten et al., 2018), suggesting the involvement of the locus coeruleus-noradrenergic (LC-NA) neuromodulatory system (Murphy et al., 2014) in value-based decision-making during learning. A prominent theory in the field, the adaptive gain theory, describes a leading role of the LC-NA system in the explore-exploit trade-off (Aston-Jones & Cohen, 2005). In a putative phasic mode, phasic LC activity responds to task-related events to optimize performance, occurring alongside a moderate level of tonic activity. In a tonic mode, tonic LC activity takes over as neurons stop responding phasically when utility in the task diminishes, leading to distractibility and poorer performance but also to the exploration of alternative options. This tonic mode may be adaptive by aiding a change in behaviour if either the current task is no longer rewarding or if the environment has changed.

Anxious individuals show reduced adaptability of learning rate to changes in volatility, in threatening (Browning et al., 2015) and rewarding contexts (Huang et al., 2017). In an experiment that posed the potential of a shock, individuals high in trait anxiety, as compared to those with lower anxiety, exhibited a blunted increase in learning rate when moving from a stable to volatile environment, i.e., trait anxiety was associated with a reduced ability to appropriately adapt to a volatile environment (Browning et al., 2015). A similar study reported elevated learning rates for negative outcomes in a volatile punishment environment in those with mood and anxiety symptomatology compared to controls (Aylward et al., 2019). A study on state anxiety, i.e., current anxious arousal as a separate (but associated) construct to trait or chronic anxiety, examined anxiety-associated differences in uncertainty representations, describing how, in a highly volatile and noisy environment, state-anxious individuals exhibit a reduced estimate of volatility, leading to a lower learning rate compared to those with less anxiety (Hein et al., 2021).

It has recently been proposed that the difference in uncertainty processing in anxious individuals, rather than being associated with volatility, is in fact due to an underlying deficit in estimating stochasticity (noise) (Piray & Daw, 2021). The authors suggest that anxiety primarily disrupts inference about stochasticity but that the learner misinterprets fluctuations due to stochasticity as a signal of change, i.e., volatility. Predictions from Piray & Daw’s model were partly based on data from Huang and colleagues (Huang et al., 2017), obtained from a change point detection task. They found that individuals with high anxiety had difficulty determining if an action was associated with an outcome by chance (noise) or by some statistical regularity in the environment (volatility). This was indicated by a higher lose-shift rate in people with high compared to low anxiety, i.e., those with high anxiety were more inclined to switch to choosing the other option after receiving negative feedback.

In the afore-mentioned studies, volatility and noise were either fluctuating in a complex, (pseudo-) random manner across the experiment (Huang et al., 2017; Hein et al., 2021), or there was a change from a stable to volatile environment (Behrens et al., 2007; Browning et al., 2015; Lawson et al. 2017; Manning et al., 2017). To the authors’ knowledge, few if any studies to date have systematically isolated changes in volatility from changes in noise at the experimental level. In the current study, we use a standard probabilistic reversal learning task to explicitly test how changes in uncertainty impact learning behaviour, by orthogonally manipulating the level of noise and volatility in the environment. We examine changes across these different levels of uncertainty and address how different combinations of noise and volatility across experimental blocks affect learning in adults with and without high levels of trait anxiety. Consistent with prior work (Yu & Dayan, 2005, Piray & Daw, 2021), we predicted that across all participants, volatility would generally increase learning rates and noise would generally decrease learning rates. However, in highly anxious individuals elevated learning rates in response to noise would support the proposal that noise is being mistaken for volatility.

## Results

Participants completed four blocks (135 trials each) of a probabilistic reversal learning task (Fig 1A), in a 2x2 orthogonal design with low or high volatility combined with low or high noise. The effects of volatility and noise were assessed by doubling the rate of volatility between low and high volatility conditions (i.e. 3 vs. 7 reversals), and by reducing the level of noise by 10% from low to high noise conditions (see Methods). Reversal structure and averaged trial-by-trial responses for each condition are shown in Fig 1B. On each trial participants had to choose between a blue and an orange cup to try and find the reward (gold coin) hidden underneath. They were instructed to try to choose the cup that gives more reward on average and across time. They were told that the best cup may change over time and they should figure out when to stop choosing one cup and switch to the other.

**Fig 1.**
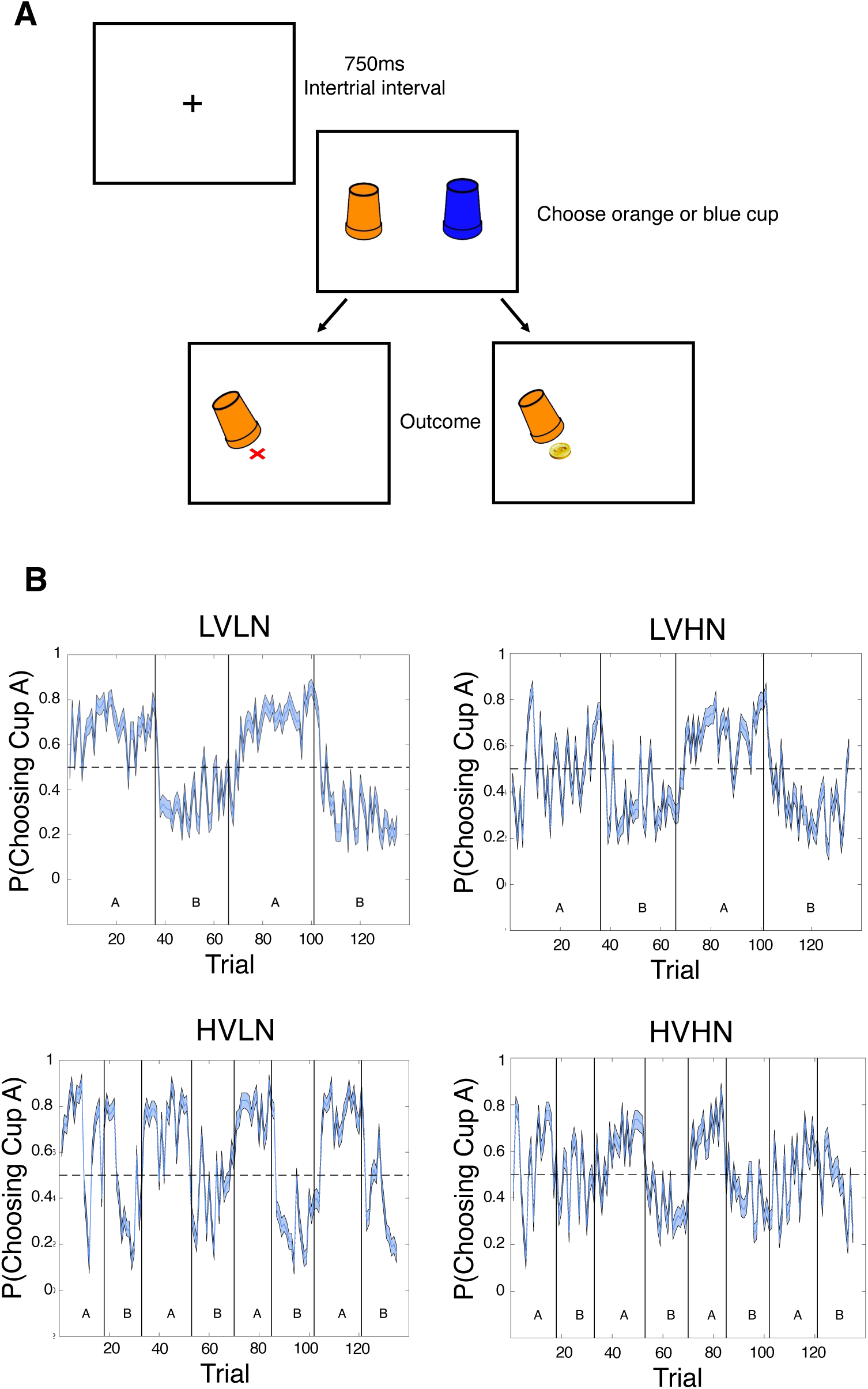
Experimental paradigm and trial-by-trial behavioural responses. **A)** On each trial, participants were presented with a blue and orange cup and had to choose one of them, receiving either a coin as positive feedback or a red ‘x’ as negative feedback. **B)** Probability of choosing one of the cups (denoted here as Cup A) across time in the full sample (N=80). Vertical lines represent reversals in contingency, with three reversals occurring in the low volatility condition and seven reversals for high volatility. Trial-by-trial responses are shown for all four experimental conditions: low volatility with low noise (LVLN), low volatility with high noise (LVHN), high volatility with low noise (HVLN), and high volatility with high noise (HVHN). Confidence intervals are ± 1 SEM around the group mean.

### Data preprocessing

As participants were not instructed to respond quickly, we set a lenient upper cut-off of 4000ms for average reaction times (RTs) to ensure they were not taking breaks within blocks of trials. Average RTs were below the predetermined cut-off of 4000ms for all participants, allowing all data to be included in further analysis (see S1 Fig for RTs in each condition). Four participants had an average RT of greater than 2 seconds, the longest of which was 3.13 seconds. Since the environment was highly changeable there was no objective cut-off for accuracy. A lower threshold for exclusion was therefore defined as less than 2.5 SD from the mean overall accuracy (62.24±8.43%), calculated as 41%. No participants surpassed this threshold; four participants scored lower than 50% accuracy, the lowest of which was 46.67%.

All participants completed the state-trait anxiety inventory (STAI) to capture the extent of anxious traits, with a mean STAI trait score of 50.75±10.21. As has been demonstrated previously (Gillan et al., 2016), the state and trait components of the STAI were highly correlated (*r*=0.71, p<.001). Participants were split into low-moderate and high anxiety groups according to their level of anxious traits (N=28 and N=27 respectively). Based on a tertile split, the cut-off for STAI trait scores was <=46 for the low-moderate anxiety group (lowest possible score was 20) and >=54.67 for the high anxiety group (highest possible score was 80). For presentation purposes, we will refer to the low-moderate anxiety group as low ANX, relative to the high ANX group.

### Demographics

The full sample (N=80) consisted of 61 female participants, 16 males, and 3 responding as neither (non-binary, other, or prefer not to say categories), with a mean age of 24.31 ± 7.67 years. After splitting data into low and high ANX groups, there were no significant group differences in age (low ANX: 24.14 ± 8.39 years, high ANX: 22.44 ± 4.41 years; W = 381.5, p = .959), gender (low ANX: 71.3% female, high ANX: 74.07% female; *X*^2^(1, 55) = 4.32, p =0.365), or education level (binarized for obtaining bachelor degree or higher; low ANX: 53.57%, high ANX: 51.85%; Fisher’s exact test: p=1).

### Behavioural analysis

#### Full sample

Average RT and accuracy across the full sample was 808.06±482.55ms and 62.24±8.43%, respectively. An analysis of task accuracy showed a significant main effect of noise level (F(1,79)=79.149, p<.001, η_p_^2^=0.500; see Table 1 and Fig 2A for behavioural results), with greater accuracy in the low compared to high noise conditions. To assess immediate responses to feedback, or reactivity, we looked at the tendency to stick with choosing the same option immediately after receiving positive feedback (win-stay) and to shift responding after negative feedback (lose-shift).

**Fig 2.**
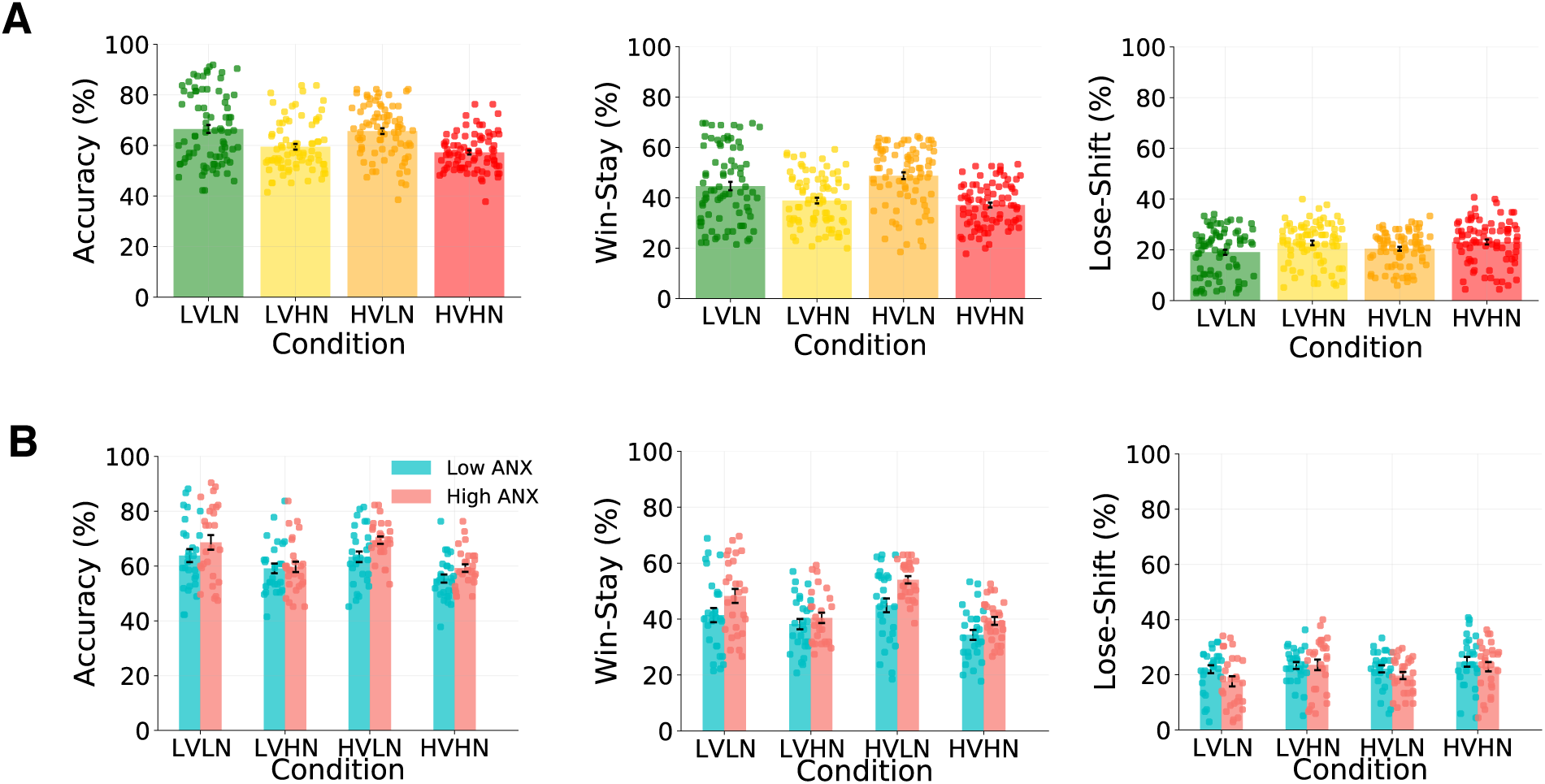
Behavioural results. Accuracy, win-stay responses, and lose-shift responses in **(A)** the full sample (N=80), and **(B)** the low (N=28) and high (N=27) ANX groups. All measures demonstrate a visible effect of noise level on behaviour, with better task performance, more win-stay behaviour, and less lose-shift behaviour under low compared to high noise conditions (all p<.001 in full sample). Participants showed more win-stay responses when moving from low to high volatility in the context of low noise, and less win-stay behaviour when moving from low to high volatility in the context of high noise (p<.001 in full sample). The high ANX group had significantly more win-stay responses than the low ANX group under low (p_bonf_=.015) but not high noise (p_bonf_ = .962). The high ANX group also showed significantly more lose-shift responses in the high compared to low noise conditions (p_bonf_ <.001) whereas the low ANX group did display any noise-related differences in lose-shift behaviour (p_bonf_=0.132). Error bars are ± 1 SEM.

**Table 1.**
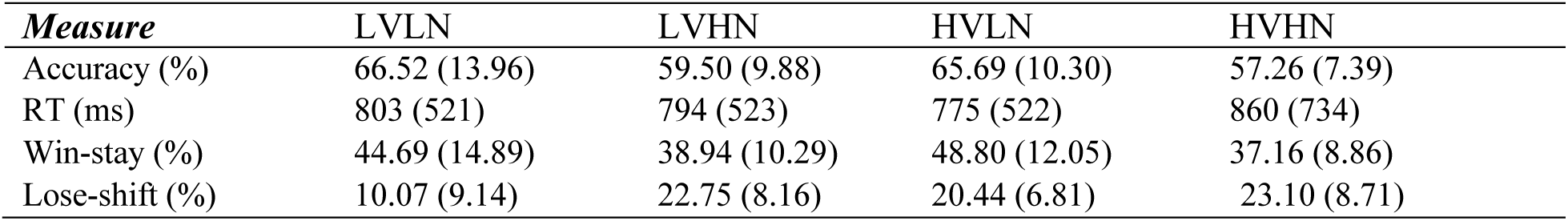
Behavioural results for the full sample. Numbers in brackets represent ±1 SD.

For win-stay behaviour, we found a significant main effect of noise (F(1,79)=114.069, p<.001, η_p_^2^=0.591), with greater win-stay responses in low noise blocks. There was also a volatility*noise interaction (F(1,79)=15.022, p<.001, η_p_^2^=0.160), with more win-stay responses when moving from low to high volatility in the context of low noise, and less win-stay behaviour when moving from low to high volatility in the context of high noise. For lose-shift behaviour we found a significant main effect of noise only (F(1,79)=35.465, p<.001, η_p_^2^=0.310).

#### Anxiety groups

Average RT and accuracy were 914.95±628.75ms and 60.42±7.80% for the low ANX group, and 721.78±244.20ms and 64.25±7.50% for the high ANX group. There was no difference in which condition participants received first (LVLN, LVHN, HVLN, or HVHN) across ANX groups (*X*^2^ (3,55) = 3.4, p=0.337). An analysis of task accuracy showed a significant main effect of noise level (F(1,53)=58.74, p<.001, η_p_^2^=0.51; see Table 2 and Fig 2B), with higher accuracy in the low compared to high noise conditions, as was observed in the full sample analysis. There were no other main effects or interactions (all p>.1). However, the best fitting model according to a Bayesian repeated-measures ANOVA included main effects of both noise and ANX group (BF=7.14, R^2^=0.521, 95% CI=[0.436, 0.595]), with the high ANX group exhibiting higher accuracy in general regardless of volatility or noise level, closely followed by a noise only model (BF=6.20, R^2^=0.514, 95% CI=[0.425, 0.587]). This suggests that both noise and ANX level played an independent role in task performance.

**Table 2.** Behavioural results for low ANX and high ANX groups. Numbers in brackets represent 1 SD.

| *Measure* | *Low ANX* |  |  |  |
| --- | --- | --- | --- | --- |
|  | LVLN | LVHN | HVLN | HVHN |
| Accuracy (%) | 63.78 (12.64) | 59.13 (9.36) | 63.36 (10.32) | 55.40 (7.85) |
| RT (ms) | 888 (516) | 939 (667) | 893 (735) | 939 (1017) |
| Win-stay (%) | 41.43 (13.52) | 38.20 (9.80) | 44.92 (13.22) | 34.37 (9.33) |
| Lose-shift (%) | 22.01 (7.48) | 23.39 (6.57) | 22.17 (6.71) | 24.68 (9.28) |

| *Measure* | *High ANX* |  |  |  |
| --- | --- | --- | --- | --- |
|  | LVLN | LVHN | HVLN | HVHN |
| Accuracy (%) | 68.64 (13.84) | 59.67 (9.88) | 69.44 (6.98) | 59.26 (7.19) |
| RT (ms) | 658 (245) | 675 (321) | 689 (322) | 865 (483) |
| Win-stay (%) | 48.29 (13.15) | 40.44 (9.49) | 54.05 (6.80) | 39.37 (7.35) |
| Lose-shift (%) | 17.67 (9.59) | 23.59 (10.01) | 19.73 (6.61) | 22.96 (8.82) |

**Table 2a.**
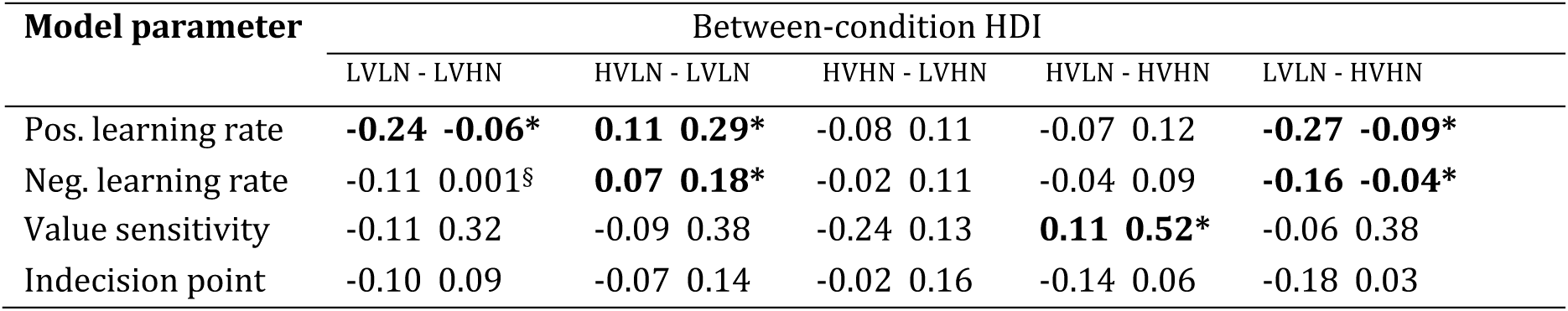
Differences in full sample model parameter posterior distributions.

Assessing win-stay behaviour, we found a significant main effect of noise (F(1,53)=94.99, p<.001, η_p_^2^=0.642), with more win-stay responses for low compared to high noise conditions. Similar to the full sample, there was a volatility*noise interaction (F(1,53)=17.43, p<.001, η_p_^2^=0.247), with more win-stay responses when moving from low to high volatility in low noise conditions, and less win-stay behaviour when moving from low to high volatility in high noise conditions. Crucially, there was a significant noise*ANX group interaction (F(1,53)=5.51, p=.023, η_p_^2^=0.094), with a larger decrease in win-stay behaviour between low and high noise conditions in the high ANX compared to low ANX group. Post-hoc comparisons showed significantly more win-stay responses for the high compared to low ANX group under low noise (M±SE=7.99±2.55, t=3.13, p_bonf_=.015) but not under high noise (M±SE=3.62±2.55, t=1.42, p_bonf_=.962). Overall, this interaction suggests a differential impact of low noise on win-stay behaviour according to level of anxiety, with highly anxious people benefitting more from less noisy conditions. Finally, there was also a main effect of ANX group, with more win-stay behaviour in general in the high compared to low ANX group (F(1,53)=5.97, p=.018, η_p_^2^=0.101). A Bayesian repeated-measures ANOVA confirms these findings on win-stay behaviour but also includes a main effect of volatility in the best fitting model, i.e., main effects of noise, ANX group, and volatility, as well as volatility*noise and noise*ANX group interactions (BF=13.491, R^2^=0.638, 95% CI=[0.574, 0.688]).

For lose-shift behaviour there was a significant main effect of noise, with more lose-shift responses in high compared to low noise conditions (F(1,53)=30.83, p<.001, η_p_^2^=0.368). There was also a significant noise*ANX group interaction (F(1,53)=5.03, p=.029, η_p_^2^=0.087), with less lose-shift behaviour for the high compared to low ANX group under low noise conditions, but similar behaviour between ANX groups under high noise conditions. Post-hoc comparisons revealed significantly more lose-shift responses in the high compared to low noise conditions in the high ANX group (M±SE=4.58±0.84, t=5.46, p_bonf_ <.001) with no meaningful difference in this measure for the low ANX group (M±SE=1.94±0.82, t=2.36, p_bonf_=0.132). These anxiety-related differences in lose-shift behaviour mirror those seen for win-stay behaviour; compared to less anxious people and in the context of low noise, those with higher anxiety stick with the same choice more after receiving positive feedback and shift to the other option less after receiving negative feedback. In line with the findings of Huang and colleagues (Huang et al., 2017), lose-shift responding was driven by greater sensitivity to the level of noise in more anxious participants. Although a Bayesian version of the analysis includes noise only as the best fitting model (BF=9.95, R^2^=0.576, 95% CI=[0.500, 0.635]), there was also evidence for a model including main effects of noise and ANX group, and a noise*ANX group interaction (BF=4.36, R^2^=0.581, 95% CI=[0.497, 0.643]), and for a model with just main effects of noise and ANX group (BF=4.36, R^2^=0.575, 95% CI=[0.489, 0.639]).

### Hierarchical Bayesian model analysis and model validation

Six learning models were fit to each condition for (1) the full sample and (2) separate high and low ANX groups, and the model with the lowest LOOIC value was selected for further analysis for each of these groups (see S1 Table and S2 Table). The winning model was the fictitious update reward-punish model with indecision point (FU-RP-IP). For the full sample, the FU-RP-IP model fit the data best in three out of four models, and for the high and low ANX groups, it was the best fit in seven out of eight models. There was one instance, the LVHN condition, where a very similar model without the indecision point was the best fit to the data. Here we present results from the FU-RP-IP model. An analysis of the model without indecision point also showed a similar pattern of results and group differences as those presented here and does not alter our conclusions (see S2 Fig and S3 Table).

Model validation of the winning ANX group models were carried out using posterior predictive checks, to ensure that the model mimics the data quite well, i.e., validate that the model can recapture actual behaviour (see S3 Fig). Furthermore, using two different simulation methods we simultaneously demonstrate both adequate parameter recovery of the ANX group models and the constraining nature of hierarchical group-level parameter distributions on lower-level individual-level parameters (see S1 Text and S4 Fig).

#### Full sample model

Median and standard deviation values of posterior distributions are presented in the S4 Table, with posterior group-level distribution plots shown in Fig 3A (see S5 Fig for indecision point plots). For all comparisons between group-level posterior distributions, we use the highest density interval (HDI), with any interval that doesn’t cross 0 indicating a meaningful difference between distributions (see Methods). The positive learning rate parameter was lowest for the LVLN condition, with significantly higher values in the HVLN (HDI = [0.11, 0.29]) and LVHN (HDI = [0.06, 0.24]) conditions, i.e., there was an increase in positive learning rate due to both volatility and noise. There was a volatility-related increase in negative learning rate from LVLN to HVLN (HDI = [0.07, 0.18]), and a noise-related increase in the value sensitivity parameter from the HVHN to HVLN condition (HDI = [0.11, 0.52]). Finally, there was a significant increase in both positive and negative learning rate when comparing the most uncertain (HVHN) to least uncertain (LVLN) conditions (HDI = [0.09, 0.27] and [0.04, 0.16] for positive and negative learning rate, respectively). There were no other significant differences in pairwise-comparisons of parameter distributions (all HDIs overlapped 0).

**Fig 3:**
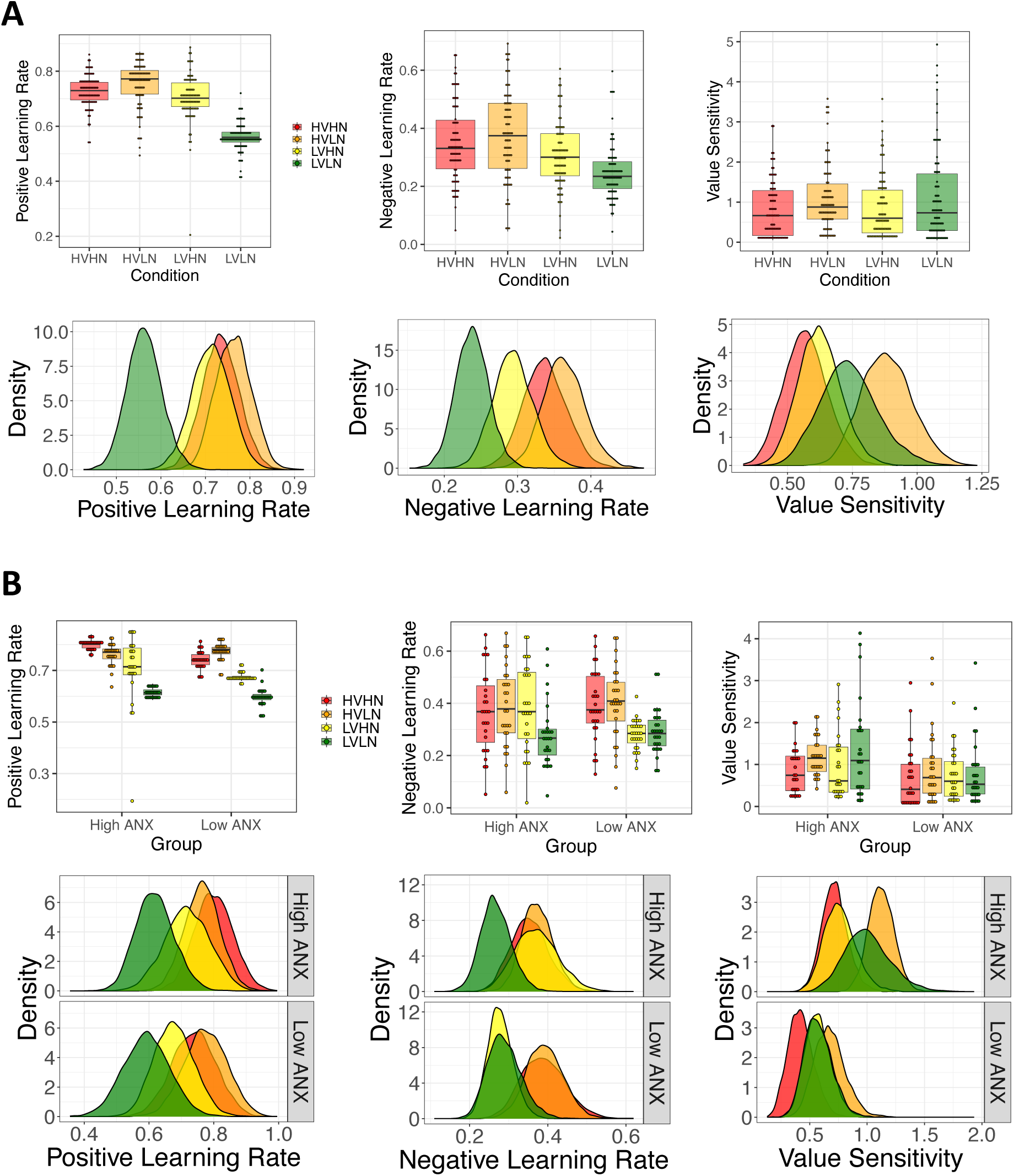
Individual-level and group-level model parameters from the winning hierarchical Bayesian model. **(A)** model parameters of the full sample (N=80), showing the median of individual-level parameters (upper panel), and group-level posterior distributions (lower panel). The LVLN condition (in green) can be seen to engender lower positive and negative learning rates in general. **(B)** model parameters for the low (N=28) and high (N=27) ANX groups. Crucially, negative learning rate for the LVHN condition (in yellow) overlaps with the LVLN condition in the low ANX group, i.e., similar distributions for high versus low noise under low volatility, but is shifted to higher values and overlaps the HVHN condition in the high ANX group. Value sensitivity distributions are higher in general for high compared to low ANX groups. Note that condition-based analyses of differences according to ANX group are carried out on the group-level distributions only.

#### Anxiety group models

Group-level posterior distribution plots are shown in Fig 3B (also see S6 Fig). Median and standard deviation values of posterior distributions of each ANX group, condition and parameter are shown in S5 Table. Comparing group-level parameters for each condition, there was a significant ANX group difference in the value sensitivity parameter for the LVLN (HDI=[0.03,0.78]), HVLN (HDI=[0.16,0.71]) and HVHN (HDI=[0.05,0.55]) conditions, with greater sensitivity for high compared to low ANX. There was no such difference in the LVHN condition (HDI=[-0.18, 0.47]. See Table 3 for all group and condition comparisons. Taken together, these findings suggest a more exploitative choice strategy in general in the high ANX group, and that choice strategy of the low and high ANX groups was most similar in the LVHN condition.

**Table 3.**
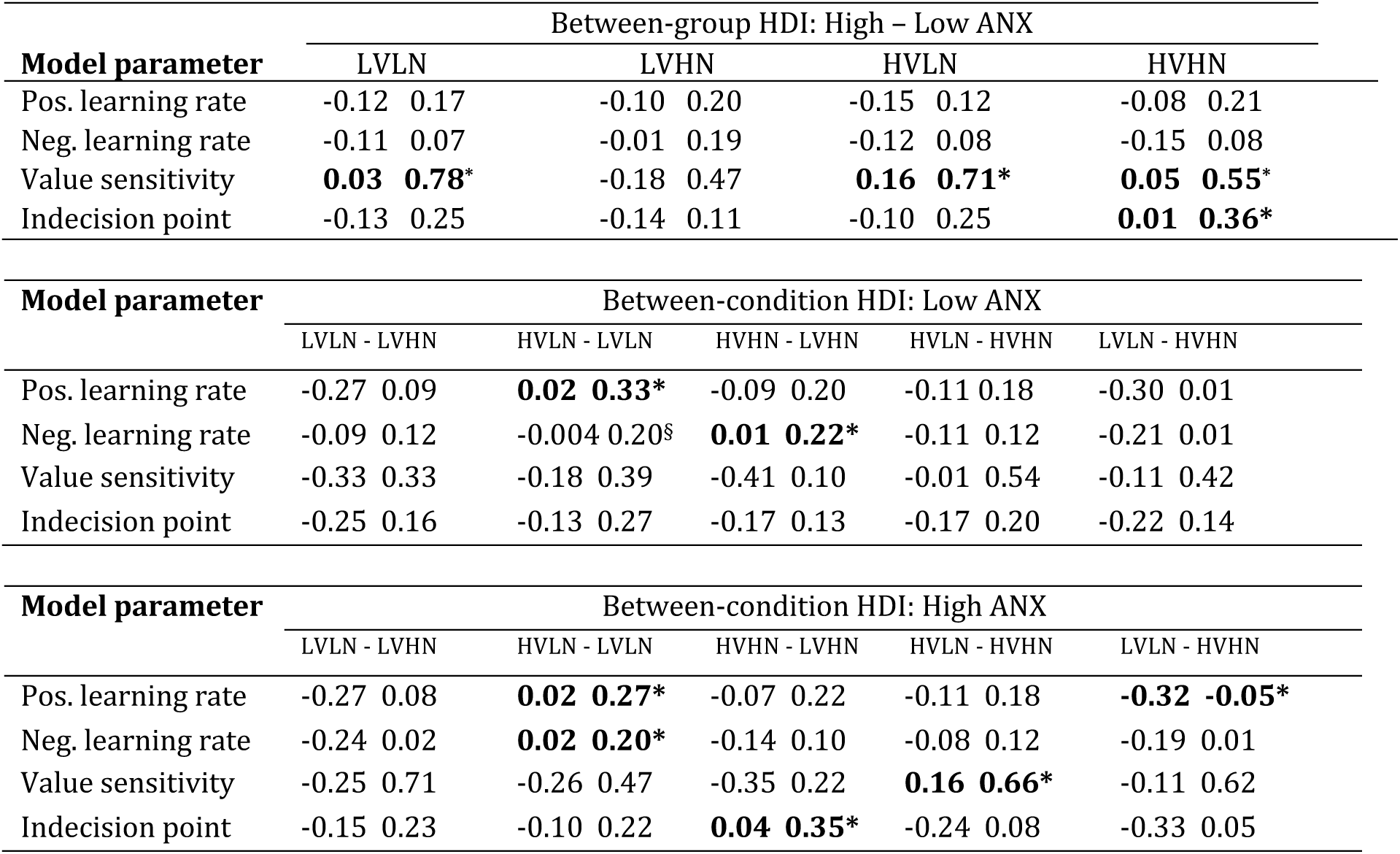
Differences in ANX groups model parameter posterior distributions. Highest density intervals (HDI) on group and condition parameter posterior distribution differences, showing the lower and upper bounds of the 89% HDI. HDI that does not overlap 0 indicates a meaningful difference between distributions. Between-group HDI per condition (top), and between-condition HDI for each group (middle: Low ANX, bottom: High ANX).

Comparing conditions in each ANX group separately, there was a significantly higher positive learning rate in the HVLN compared to LVLN condition in both low and high ANX groups (HDI=[0.02,0.33] and [0.02,0.27], respectively), i.e. under low noise, a shift from low to high volatility was associated with an increase in positive learning rate regardless of anxiety level, reflecting findings from the full sample model. For the same HVLN versus LVLN comparison, there was also a significant shift in negative learning rate in the high ANX group (HDI=[0.02,0.20]).

Although not significant, the low ANX group showed a very similar shift (see Table 3). Notably, the low ANX group showed a significantly higher negative learning rate in the HVHN compared to LVHN condition (HDI=[0.01,0.22]), but this difference was not exhibited in the high ANX group, with the HVHN and LVHN distributions largely overlapping (HDI=[-0.14,0.10]; see Fig 3B). This suggests that highly anxious people employ a similarly high negative learning rate under both high noise conditions, without making an adjustment for the change in volatility. Finally, the high ANX group showed significantly higher value sensitivity in the HVLN compared to HVHN condition (HDI=[0.16,0.66]), i.e., under high volatility, those with high ANX showed substantially more exploitation of learned value differences for low compared to high noise.

#### Relationship between anxious traits and learning in the full sample

Based on the reported ANX group findings, we assessed whether there was a relationship between anxious traits in the full sample and 1) win-stay behaviour, 2) lose-shift behaviour, and 3) negative learning rate (see Fig 4). There was a significant positive relationship between overall win-stay behaviour and STAI traits, regardless of task condition (r=.261, p=.019). Higher anxious traits were also associated with more win-stay behaviour in low compared to high noise conditions, although this did not pass the significance threshold (r=0.210, p=.061). For lose-shift behaviour, the reported interaction between low/high ANX group and noise level was corroborated using the distribution of anxious traits in the full sample; those with higher anxious traits displayed significantly more lose-shift behaviour in high compared to low noise conditions (r=.258, p=.021). Examining low and high volatility conditions separately, this correlation appeared to be strongly driven by the low volatility condition (LVHN – LVLN; r=.292, p=.009), i.e., noise-related lose-shift behaviour scaled with levels of anxious traits under low but not high volatility. Although not passing the significance threshold, we also found an indication of higher negative learning rates in the LVHN compared to LVLN condition with increasing anxious traits (r=0.20, p=.069).

**Fig 4.**
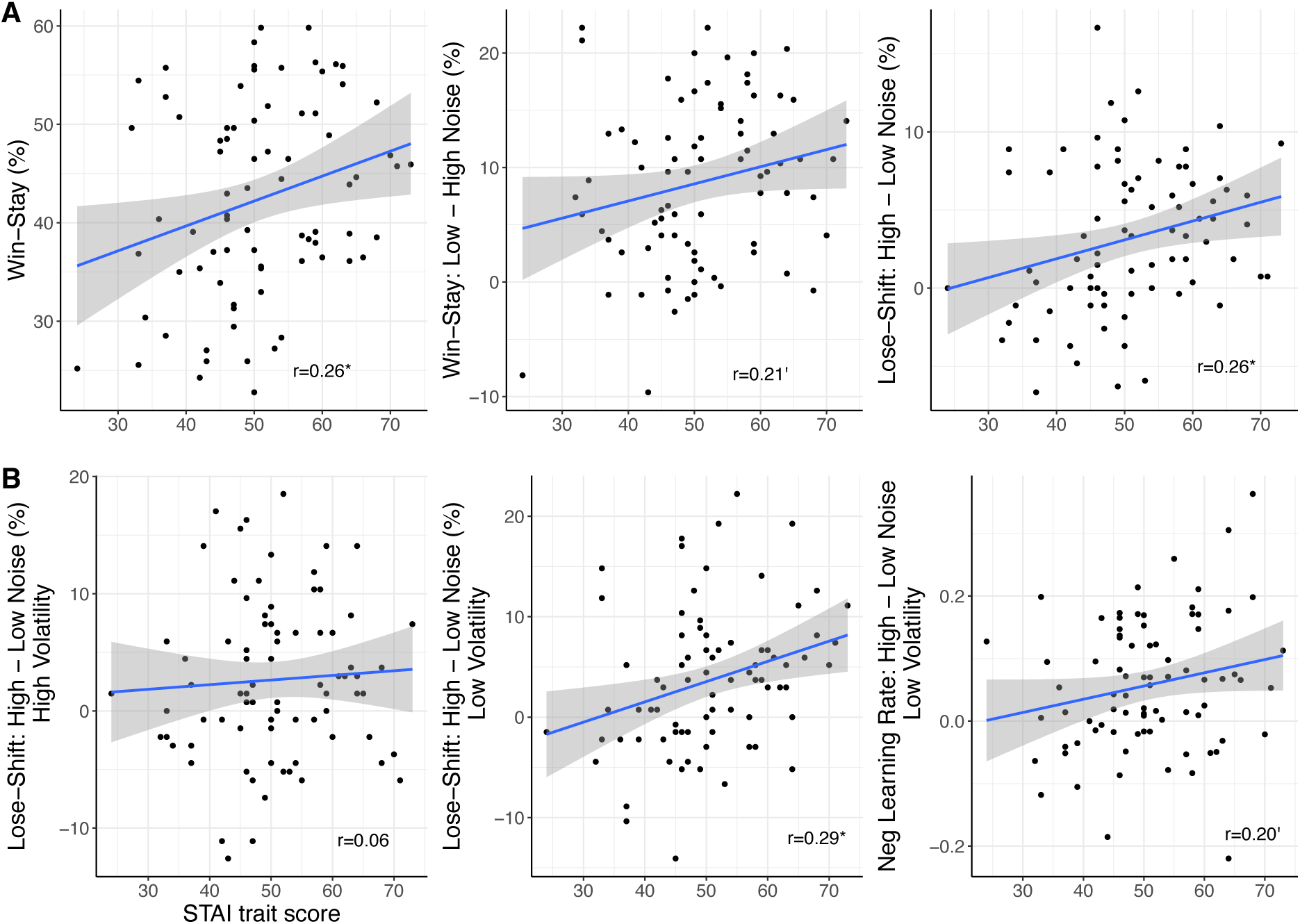
Relationship between learning and STAI traits in full sample (N=80), extending ANX group-based findings. **A)** Significant positive correlation between anxious traits and general win-stay behaviour (p=.019; left panel); a positive, trending-only relationship between anxious traits and win-stay behaviour in low compared to high noise conditions (p=.061; middle panel); and a significant positive relationship between anxious traits and lose-shift responses in high compared to low noise conditions (p=.021; right panel). **B)** No relationship between anxious traits and lose-shift behaviour in high compared to low noise conditions under high volatility (p<.1; left panel), but crucially, a significant positive correlation between anxious traits and lose-shift behaviour in high compared to low noise conditions under low volatility (LVHN – LVLN, p=.009; middle panel); and a positive, trending-only relationship between anxious traits and negative learning rate for the same comparison (LVHN – LVLN, p=.069; right panel). Grey bands represent 95% confidence intervals.

Previous research that suggests a relationship between task accuracy and the value sensitivity parameter (Jahfari et al., 2019; McCoy et al., 2019), and a relationship between lose-shift behaviour and negative learning rate (St. Onge et al., 2011). In supplementary analyses (S7 Fig) we also confirm a highly significant association between noise-related changes in accuracy and noise-related changes in value sensitivity, regardless of volatility level (r=.751, p<.001), as well as a tight relationship between noise-related changes in lose-shift behaviour and noise-related changes in negative learning rate under low volatility (r=.487, p<.001). Taken together, these associations based on the full sample largely confirm those found in the ANX group-level analyses and extend those findings to demonstrate the effect of anxious traits and noise on learning model parameters.

## Discussion

Adapting to different types of uncertainty is a daily endeavour, however, few studies have explicitly teased apart the two main sources of uncertainty – stochasticity of the incoming information (noise) and the changeability of the signal over time (volatility), in both health and disorder. People with anxiety disorders are known to experience difficulties with tolerating uncertainty (Bishop 2007; Carleton, 2016), making this an important avenue of research for the effective treatment of the negative effects of anxiety. In this online study, participants with a range of anxious traits attempted to learn which stimulus was optimal across time, in a fully orthogonal RL task with low or high volatility combined with low or high noise.

We found that participants performed better under low compared to high noise conditions, regardless of volatility level. Remarkably, even a 10% difference in noise level between conditions appeared to have a greater influence on behaviour than doubling the level of volatility. We also found more win-stay and less lose-shift behaviour under low versus high noise conditions. With regards to anxiety, there were more win-stay responses in general for the high compared to low ANX groups, and win-stay behaviour scaled with increased levels of anxious traits in the full sample. This is a novel finding, as previous literature suggests that only lose-shift behaviour is inclined to be affected by anxiety levels (Huang et al. 2017). There was also a greater drop in win-stay responses and a greater increase in lose-shift responses when moving from low to high noise conditions for the high compared to low ANX groups. Results from the full sample indicate that people with higher anxious traits are more sensitive to differences in noise levels, with significantly less lose-shift behaviour under low compared to high noise conditions, specifically in the context of less volatility. Overall, our results point towards noise having a more influential impact on behaviour than volatility, both in the general population and in those with heightened anxiety especially.

Across all participants, we demonstrate that in the less noisy conditions (75:25 reward contingency), there is a higher positive and negative learning rate for high compared to low volatility, corroborating previous studies showing an increase in learning rate from a stable to volatile environment (Behrens et al., 2007; Browning et al., 2015; Lawson et al., 2017; Manning et al., 2017). This was not the case, however, for higher noise conditions (65:35 reward contingency), with high and low volatility learning rate distributions largely overlapping. Taken alongside behavioural findings showing increased win-stay responses for high versus low volatility under low noise but not high noise conditions, these results indicate that combining different levels of noise and volatility substantially influences learning from feedback. For positive learning rate, there were no ANX group differences, and both low and high ANX groups demonstrated the significant increase in positive learning rate from low to high volatility under low noise. Under high volatility, there was greater value sensitivity for low compared to high noise in the full sample. This indicates that people behave in a more exploitative manner, using their knowledge of learned value differences between options, in situations that are less noisy, particularly when the environment is more volatile. We furthermore found greater value sensitivity in the high compared to low ANX group in general during this dynamic learning task.

Predictions from Piray and Daw (Piray & Daw, 2021) suggest that anxiety primarily affects inference about noise (stochasticity) but that the learner misinterprets fluctuations due to noise as a change caused by volatility. The authors present a model demonstrating greater change in a volatility parameter and a flatlined response (lesion) to stochasticity in anxious people compared to controls. In the current task, the LVHN condition appears to be the only context where both low and high ANX groups perform behaviourally similar, with comparable accuracies and lose-shift behaviour. In the full sample, we found more lose-shift behaviour in LVHN compared to LVLN for those with higher anxious traits. This noise-related change in lose-shift behaviour is associated with anxious traits only under low and not high volatility. Compared to the LVLN “baseline” condition, the high ANX group shows an increase in negative learning rate in LVHN (although not significant with these contingencies), leading to a completely overlapping distribution with the HVHN condition, whereas the low ANX group shows a significant difference between the LVHN and HVHN conditions. In conjunction with these changes in negative learning rate, the significant difference in value sensitivity between the low and high ANX groups seen in all other conditions is not present in the LVHN condition. Together, these findings offer support for Piray & Daw (2021), suggesting that perhaps people with high anxiety perceive the high noise occurring with low volatility as being caused by an increase in volatility.

A prevalent RL paradigm, the probabilistic selection task (PST), involves learning about three stimulus pair options, with stable reward contingencies 80:20, 70:30 and 60:40 for each of the pairs, denoted as AB, CD, and EF, respectively (Frank et al., 2007). Its use of three stimulus pairs rather than one distinguishes it from the task used here but it is nonetheless informative in understanding how differences in noise levels across different pairs of stimuli might affect performance. Many studies using the PST have confirmed increased task performance for the more certain (least noisy) AB pair relative to the other pairs (Jahfari et al., 2019; van Slooten et al., 2018; McCoy et al. 2019). The value sensitivity parameter has also previously been associated with overall performance in the PST (Jahfari et al., 2019, McCoy et al., 2019). Here, we replicate this finding and extend it to changes in noise, showing a strong correlation between increases in accuracy for high compared to low noise conditions and an increase in the value sensitivity parameter for high compared to low noise.

Simulations of optimal task performance in a standard probabilistic reversal learning task indicate that value sensitivity is optimal at its maximum value in most 75:25 reward contingency environments, i.e., entirely exploitative, except for extremely volatile environments where the reward contingency reverses every ∼10 trials (Zhang et al., 2020). Although human learners tend not to perform in an optimal manner, these simulations provide a benchmark against which humans can be compared. In the current task, the highly volatile condition includes reversals every 15-20 trials. Taken alongside findings from a recent study demonstrating a reduction in inverse temperature (more exploration) in a volatile compared to stable environment (Simoens et al., 2024), and simulations demonstrating more exploratory and flexible behaviour in highly volatile environments (Zhang et al., 2020), the current findings potentially indicate that, despite the increase in task accuracy, the tendency of those with higher anxiety to exploit value differences may be a misaligned or overly rigid response to this dynamic, uncertain environment. Alternatively, or coinciding with this, anxious people may hold prior beliefs that the world is highly changeable and uncertain, leading to more optimized behavioural strategies in the face of such an environment. Future studies would need to examine the implications of this exploitative strategy under high uncertainty in terms of how it may relate to specific symptoms of anxiety such as, for example, a need for control of one’s environment.

Both dopamine and noradrenaline are known to play a shared role in learning and are functionally linked, with dopamine acting as the direct precursor to noradrenaline (Musacchio, 2013). The crucial role of dopamine during learning has been extensively studied, with activation of midbrain dopamine neurons acting as a learning signal to the striatum and other brain regions to gradually make better predictions about when rewards are likely to occur (Montague et al., 1996, 2004; Schultz et al. 1997). Prediction errors, the difference between the outcome that was expected and the actual outcome, are tracked by dopamine neurons, and are weighted by the learning rate parameters of reinforcement models. In the current study, the volatility-related differences in positive and negative learning rates under low noise likely reflect changes in dopaminergic firing due to the change in environmental volatility, with previous studies implicating the dopaminergic system in reversal learning (Cools et al., 2009; van den Bosch et al., 2022). A recent in-vivo imaging study in humans confirmed dopamine release in the striatum as a crucial component of reversal learning (Grill et al., 2024). The anxiety-related difference in negative learning rate for the LVHN condition, with a significant difference compared to the HVHN condition in the low anxiety group but overlapping distributions in the high anxiety group, potentially reflects an anxiety-related difference in dopaminergic signalling in response to feedback that depends on the underlying nature of the uncertainty. Similarly, in the full sample we present a noise-related increase in lose-shift behaviour and negative learning rate in those with high anxiety when moving from low to high noise under low volatility, without the same shift apparent in less anxious people.

Noradrenaline has previously been proposed to control the randomness in action selection, i.e., the inverse temperature/value sensitivity parameter (Doya et al., 2002). Research using the PST has shown that people who are sensitive to small value differences and make more exploitative choices have stronger choice-locked pupil dilation, while also being more accurate in their choices (van Slooten et al., 2018). Several studies have also suggested an association between pupil dilation and the noradrenergic neurotransmitter system (Murphy et al., 2014; Varazzani et al., 2015). A pharmacological study looking at the effects of attenuated noradrenergic neurotransmission on learning under uncertainty, using the β-adrenergic receptor antagonist propranolol, found that contingency learning rates were generally higher in high state anxious individuals, but this effect was diminished under propranolol (Lawson et al., 2020). The computational model used in that study, the hierarchical gaussian filter (Mathys et al., 2011), is architecturally different from the model used the current study and doesn’t involve a value sensitivity parameter. However, propranolol (i.e., diminishing noradrenergic neurotransmission) was also found to reduce the impact of prediction errors and volatility on pupil size. A mechanistic account, the adaptive gain theory, details an inverted U-shaped model to describe the relationship between LC tonic activity and flexible task switching, with a phasic mode involving moderate LC tonic activity that is associated with appropriate, task-related, exploitative phasic responding, and a tonic mode where tonic activity increases and takes over, leading to greater distractibility and the pursuit of alternative actions or choices (Aston-Jones & Cohen, 2005; Kane et al., 2017). Taken together, the overall heightened value sensitivity and better task performance seen for the high relative to low ANX group in the current study potentially suggests more phasic responding in those with high ANX under these changeable conditions, with the low ANX group at the upper, tonic LC mode of the U-shaped curve, allowing for greater flexibility to explore other options. Whether this is reflective of highly anxious people being predisposed to responding to change and expectant of it, a difference in where highly anxious people sit on the adaptive gain theory U-shaped curve, and/or a difference in how people with low and high anxiety shift along that curve in response to shifting environmental contexts, is an open avenue for future research.

Findings from this study should be interpreted in the context of certain limitations. Firstly, our task design considered just two levels of volatility and noise; reversals occurred every 30-40 trials in low volatility and every 15-20 trials in high volatility conditions, with outcome reward contingencies of 75:25 and 65:35 for low and high noise, respectively. Future studies could include more levels of uncertainty to obtain a more fine-grained picture of how learning changes across different volatility and noise combinations and according to anxious traits. For example, in the LVHN condition there was an anxiety-related shift in negative learning rate which may be more prominent if comparing across noise levels that are more distinct from each other. Secondly, separate hierarchical RL models were fit across the full sample, low ANX group, and high ANX group. In hierarchical models, low-level parameter estimates are pulled more closely together than if there were no higher-level distributions, a property termed “shrinkage” (Krushche, 2015). As is also visible from our parameter recovery simulations, the use of individual-level parameters from these models should be interpreted with caution. In the full sample model, people with the full range of anxious traits contribute to the group-level parameters, which means that individuals with low and high anxiety are all pulled closer to the group mode. This potentially obfuscates any anxiety-related differences in individual-level parameters. Finally, in terms of task design, our experimental manipulations were in a general context of high uncertainty - a highly dynamic environment - and were not compared to stable or deterministic settings. Future studies might disentangle anxiety-related differences in stable or deterministic contexts from those involving gradual increases in volatility and/or noise.

To conclude, we replicate previous findings demonstrating increased learning rates for increases in volatility, specifically in low noise conditions. Highly anxious people showed greater sensitivity to value differences in general, indicating a more exploitative strategy in highly changeable contexts, that is potentially maladaptive due to lack of flexibility. Noise, or variability in reward outcomes, appeared to play a more substantial role than volatility in behaviour with regards to anxious traits, with highly anxious people showing more sensitivity to changes in noise, indicated by a bigger shift in lose-shift behaviour and negative learning rate with changing noise level, specifically in less volatile situations. These findings suggest that highly anxious people have greater sensitivity to negative feedback when noise increases and, when applied to noisy situations encountered in daily life, this increased sensitivity likely plays a role in the negative, long-term impact of anxiety on those who suffer from it. Our study corroborates the general suggestion stemming from RL theory: if surprising outcomes are caused by high noise, particularly under low volatility as indicated here, then current actions should be based on the average over the outcomes of many previous actions. Future research would need to further probe the mechanism by which anxious people mistake increases in noise for volatility, how it might relate to specific symptoms of anxiety, and explore methods or strategies that might help anxious people to decipher changes in noise from volatility.

## Materials and Methods

### Participants

The online Gorilla Experiment Builder platform was used to develop the RL task and host this study. Eighty participants were recruited via Prolific and were automatically redirected to Gorilla using an external URL. Inclusion criteria were that participants had to use a desktop computer (not a tablet or mobile phone), were in the age range 18-65 years, and could be from any country but had to be fluent in English. The full experiment was expected to take approximately 45 minutes to complete, for which participants were compensated £6. Participants first read through an information sheet and provided informed consent by ticking their agreement to provided statements. Ethical approval for the study was granted by the Cambridge Psychology Research Ethics Committee (CPREC PRE.2019.110). Participants first completed some demographic and symptoms questionnaire data, followed by the RL task.

### Experimental paradigm

Participants performed a standard probabilistic reversal learning task, choosing between a blue and orange cup on each trial. The locations of the cups were counterbalanced across trials between the left and right side of a central fixation cross. There were four blocks with a short break between blocks. The blocks were: low volatility and low noise (LVLN), low volatility and high noise (LVHN), high volatility and low noise (HVLN) and high volatility and high noise (HVHN). The study consisted of 540 trials in total, with 135 trials in each of four blocks. The average reward contingencies (noise levels) in the low and high noise conditions, regardless of volatility, were 75:35 and 65:35 respectively. Reversals in contingency occurred every 30-40 trials for the low volatility conditions and every 15-20 trials in the high volatility conditions. To ensure that results would not be due to a specific block order, half of participants completed the low volatility conditions first and the other half of participants completed the high volatility conditions first. Low and high noise blocks were also counterbalanced within each of these groups. Participants were instructed before each block to keep trying to choose the better cup, and that the better cup may change across time. All participants were given the same sequence of trials, e.g., on a particular trial, choosing the orange cup led to the same outcome across participants. The outcome of each cup on any specific trial was always the inverse of the outcome of the other cup, i.e., if a participant chose the orange cup on trial 10 then they would have received a positive outcome (gold coin), whereas if they chose the blue cup on that trial, they would have received a negative outcome (red cross).

### Analysis of task behaviour

Behavioural performance was determined using measures of accuracy, win-stay responses, and lose-shift responses. In each block condition (LVLN, LVHN, HVLN, HVHN), one particular cup that was considered correct for each mini-block, i.e., between each reversal, regardless of whether there was a low or high noise associated with the reward contingency for that stimulus. Accuracy was calculated as the percentage of correct responses, collapsed across all mini-blocks per condition. Win-stay and lose-shift behaviour was assessed by calculating the number of times a participant stayed with choosing the same cup that was rewarded on the previous trial out of the total number of win outcomes on that condition (win-stay percentage), and the number of times a participant switched to choosing the other cup after receiving a loss for the original cup on the previous trial out of the total number of loss outcomes on that condition (lose-switch percentage). Since this was a learning and decision-making task with no instructions given to participants about how fast they should respond, and for which computational models were fit to choices and feedback only, reaction times were not of concern. This in line with previous studies using similar tasks (Crawley et al., 2020; Cools et al., 2002; den Ouden et al., 2013; Chamberlain et al., 2006). For all analyses on task accuracy, win-stay and lose-shift behaviour, we ran repeated-measures ANOVA in JASP (JASP Team, 2020), with volatility and noise levels as within-subject factors, and where relevant, ANX group as a between-subjects factor. Where appropriate, these analyses were followed up by Bayesian repeated-measures ANOVA to highlight best fitting models and other models that show moderate or strong evidence, indicated by a Bayes Factor (BF) greater than 3 or 10, respectively.

### Reinforcement learning models

In both experiments, we compared six different hierarchical Bayesian RL models using the hBayesDM package version 1.0.2 in R (Ahn et al., 2017). Each model was fit separately per task condition and participant group.

#### Model 1: Rescorla-Wagner (RW) two-armed bandit

All models incorporated the Rescorla-Wagner update rule (Rescorla & Wagner, 1972); on a given trial *t*, the value of a chosen stimulus, V_c,t_, was updated based on a prediction error, i.e., the difference between the expected value of the chosen stimulus, V_c,t-1_, and the actual outcome received, O _t-1_.

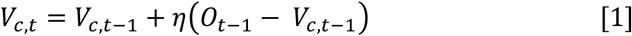

The learning rate, 𝜂 (0 < 𝜂 < 1), is a weighting on the prediction error that determines how much emphasis is placed on the current trial when updating the expected value of the stimulus.

All models also included a softmax choice function, an inverse logit transform, which relies on the difference in value between the two options to estimate the probability of choosing one stimulus over the other on a given trial.

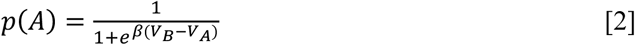

The inverse temperature parameter, β (0 < β< 10), is a weighing on the difference in value between options. We refer to the β parameter as value sensitivity, as it represents the extent to which a difference in stimulus values (obtained through an individual’s learning) determines choice, e.g., a higher β value is associated with a greater emphasis on the difference in stimulus values, whereas a low β value denotes more stochastic responses.

#### Model 2: Reward-Punish (RP)

The RP model is an expansion of the standard RW model, for which Equation 1 is split into two equations; one that is updated after receiving a reward, with a positive learning rate η_pos_, and one that is updated after receiving a punishment, with a negative learning rate η_neg_.

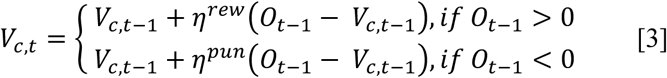

#### Model 3: Experience-weighted attraction (EWA)

The EWA model is used to account for a growing insensitivity to new information. In standard Rescorla-Wagner learning models, predictions are driven by the most recent experiences. With this model, which is extended in the hBayesDM package based on den Ouden et al. (den Ouden et al., 2013), an experience-weight parameter, called the experience decay factor ρ, denotes the importance of past experience relative to new information as the task progresses. The experience weight of the chosen stimulus on the current trial, n_c,t_, is given by:

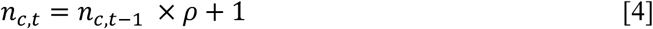

The value of the chosen stimulus on the current trial is then updated according to these experience weights, as well as a decay factor for previous payoffs φ, according to the following:

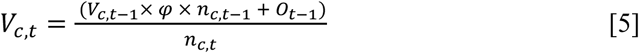

For ρ=0, trial-by-trial updates are driven by the most recent experiences, but as ρ increases to 1 all trials contribute equally to the experience weight, and thus to updates in expected value of stimuli.

#### Model 4: Fictitious Update (FU)

This model not only updates values for the chosen option (V_c,t_), but also for the option that was not chosen (V_nc,t_). This is a potential strategy because of the reciprocal relationship between the options; if the outcome of choosing one option is a win, then the outcome of having chosen the other option would have been a loss. Thus, on each trial the chosen and unchosen stimuli are updated according to the following:

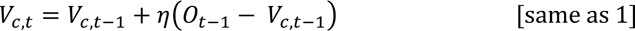

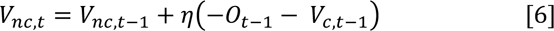

The negative sign in front of the outcome *O*_t-1_ captures the reciprocal relationship between options. The learning rate, 𝜂, is the same across both stimuli.

#### Model 5: Fictitious Update + Reward-Punish (FU-RP)

This model is a combination of both the FU and RP models, i.e., including separate reward and punishment learning rates along with value updates for both chosen and not chosen options.

#### Model 6: Fictitious Update + Reward-Punish + Indecision Point (FU-RP-IP)

This final model is the only one where an adjustment to the softmax equation (equation 2) is made, as follows:

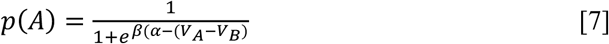

The indecision point, 𝛼, is the midpoint of the sigmoid function, i.e., where both options are equally likely to be selected. If this is far from 0, then there is a preference for one option over the other.

### Model fitting and validation

The hBayesDM package provides an accessible implementation of hierarchical Bayesian parameter estimation. In hierarchical models, group and individual parameter distributions are fit simultaneously, thereby mutually constraining and informing each other, resulting in greater statistical power over non-hierarchical methods (Ahn et al., 2011, 2017; Wiecki et al., 2013; Kruschke, 2015). Model posteriors were estimated using Markov Chain Monte Carlo (MCMC) inference. For both experiments, we fit three chains with 4,000 samples for each of the models, discarding the first 2,000 samples as burn-in. The remaining samples were used in the estimation of posterior distributions of model parameters. Convergence of the chains was confirmed using manual examination of the trace plots (hairy caterpillars, easily moving around the parameter space) and evaluation of 𝑟Bstatistics, which were all <1.1 (Gelman & Rubin, 1992). Model fits were compared using the leave-one-out cross-validation information criterion (LOOIC), to estimate out-of-sample prediction accuracy for each model using the log-likelihood of the posterior simulations of parameter values (Vehtari et al., 2017). In each experiment, the best fitting model with the lowest LOOIC value was used for further analyses. Group-level posterior estimates of parameters were compared per task condition and participant group, and the median of individual-level parameter estimates were associated with levels of clinical symptoms on questionnaires. For group-level comparisons, we use the 89% highest density interval (HDI), a credible interval (CI), on the difference between the posterior distributions. CIs computed with 89% intervals are deemed to be more stable than 95% intervals (Kruschke, 2015), particularly when using the default of 4000 samples in hBayesDM. Bayesian statisticians remind us that these CIs are arbitrary and the use of e.g., 95%, is based on frequentist statistics (McElreath, 2020). If the HDI doesn’t overlap zero, it implies that there is a significant difference between the posterior distributions.

### Symptom measures

Levels of anxious traits were measured using the state-trait anxiety inventory (STAI) (Spielberger, 1983). This includes 40 questions in total, with 20 covering how a person feels *right now* (state) and 20 relating to how a person feels *in general* (trait). This also uses a 4-point Likert scale, ranging from *not at all* to *very much so* responses. For this study we use trait scores for further analyses, as this is likely the most robust estimate and is the standard approach in similar research (Gillan et al. 2016; Aylward et al., 2019). The STAI-trait subscale has a minimum score of 20 and a maximum of 80, which may be classified as “no or low anxiety” (20-37), “moderate anxiety” (38-44), and “high anxiety” (45-80).

## Funding

RPL is supported by a Wellcome Trust Royal Society Henry Dale Fellowship [206691/Z/17/Z], an Autistica Future Leaders Award [ID: 7265], and is a Lister Institute Prize Fellow. The funders had no role in the design of the study; in the collection, analyses, or interpretation of data; in the writing of the manuscript, or in the decision to publish the results.

## Author Contributions

BM conceptualized the study and methodology, carried out data collection, curation and analysis, project administration, validation, visualization, and writing of the original draft, review & editing. RPL acquired funding, contributed to methodology, provided supervision, and reviewed and edited the manuscript.

## Data Availability Statement

Data and analysis scripts will be made publicly available upon publication of the manuscript.

## Supplementary Materials

### Supplemental Text

#### S1 Text. Parameter Recovery

Parameter recovery was performed to validate the parameter estimates and ensure that the model’s parameters are identifiable (S4 Fig). Since the same model (FU-RP-IP) was applied to all conditions, and the general features were similar across conditions, e.g., the same number of trials and a probabilistic feedback structure with several reversals in reward contingency, here we demonstrate parameter recovery on just one of the conditions – LVLN. For each ANX group, we first generated synthetic (simulated) datasets (100 simulations per participant) from the model using the observed (true) individual-level parameters from the fitted model. We then fit the FU-RP-IP model to the simulated data in two ways: 1) including all simulated data in one large hierarchical model per group, resulting in 2800 and 2700 fake datasets (subjects) in the low and high ANX group models, respectively, and 2) fitting separate hierarchical models per original participant, e.g., 100 fake participants per model. The medians of the estimated parameters were then compared to the known generating (original) parameters. A strong correlation between the simulated and true values is indicative of good parameter recovery.

For the first approach, using one large hierarchical model across all simulated datasets per group (see S4A Fig), we found that individual-level parameters were, in fact, not recovered well (all r<0.4). Inspecting this more closely, we found extremely precise group-level posterior distributions. These simulated group-level posteriors appreciably capture the medians of the original group-level posteriors (low ANX: 0.59 (±0.07) vs. 0.60 (±0.01), 0.29 (±0.04) vs. 0.26 (±0.00), 0.58 (±0.12) vs. 0.55 (±0.01), −0.05 (±0.09) vs. −0.06 (±0.01) for true versus simulated positive learning rate, negative learning rate, value sensitivity and indecision point parameter estimates, respectively; high ANX: 0.62 (±0.06) vs. 0.61 (±0.00), 0.27 (±0.04) vs. 0.25 (±0.00), 1.00 (±0.20) vs. 0.97 (±0.02), 0.01 (±0.08) vs. −0.02 (0.01) for the same parameter comparisons). Given the large number of fake participants in the simulated models (2700 and 2800 for low and high ANX groups), we interpret the non-recoverability at the individual-level to be due to the precision of these group-level distributions. Group-level parameters are known to pull individual-level parameters more tightly together, a process called “shrinkage” (Kruschke, 2015), likely leading to the simulated and true parameters effectively being uncorrelated.

With the second approach, fitting separate hierarchical models per original participant, we found that individual-level parameters were recoverable (see S4B Fig), with very strong correlations for negative learning rate (r=0.965, p<.001), value sensitivity (r=0.999, p<.001) and indecision point parameters (r=0.934, p<.001), and a lesser, but moderate correlation for positive learning rate (r=0.582, p<.001). Plotting the original group-level distributions beneath the simulated individual estimates, we show that the simulated individual-level positive learning rate estimates that deviate most from the true values fall within the range of the true group-level posterior distribution, i.e., in the true fitted model, the group-level estimates constrain the individual-level parameters, whereas individual-level estimates from this simulation method that only includes datasets from the same original participant, are not constrained in the same way. The relatively wider, less precise true group-level distribution for positive learning rate, compared to the other parameter distributions, is also indicative of greater variability at the individual level, which is likely what we are capturing with the results of this simulation method.

Together, these parameter recovery methods indicate that the set of estimated parameters describes the observed data well at the individual-level and particularly well at the group-level, which is the basis for our ANX group analyses.

#### Supplemental Figures

**S1 Fig.**
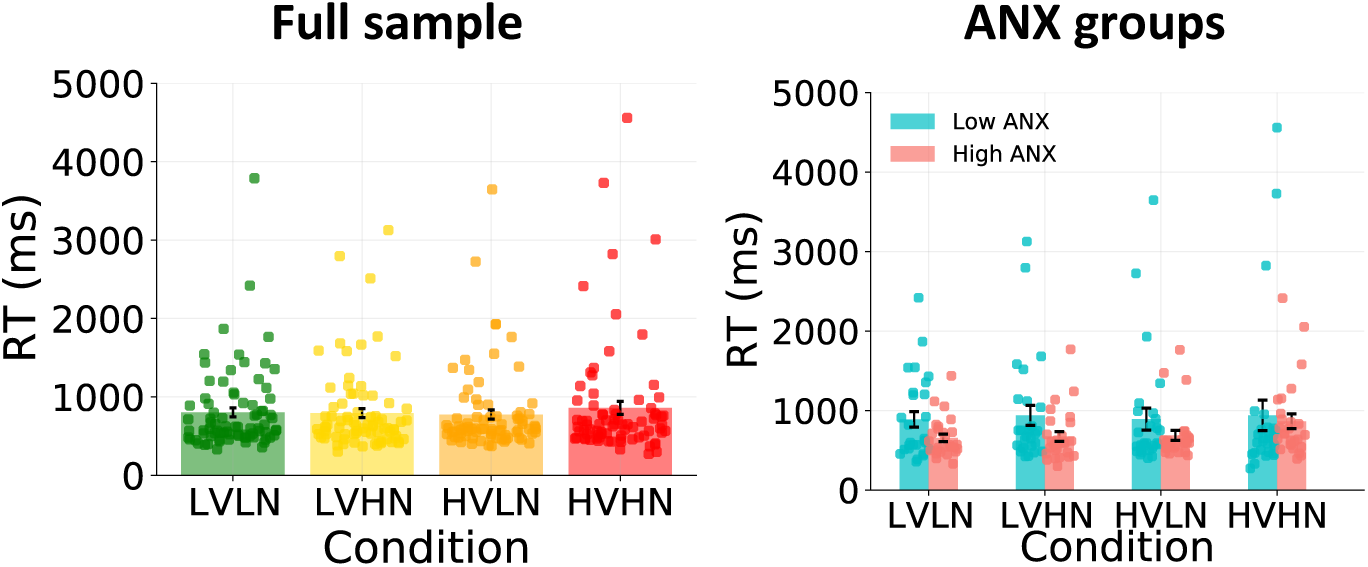
Reaction times. A repeated-measures ANOVA on RTs showed no main or interactive effects of volatility or noise, and no interaction between anxiety groups and either of these conditions (all p>.1).

**S2 Fig.**
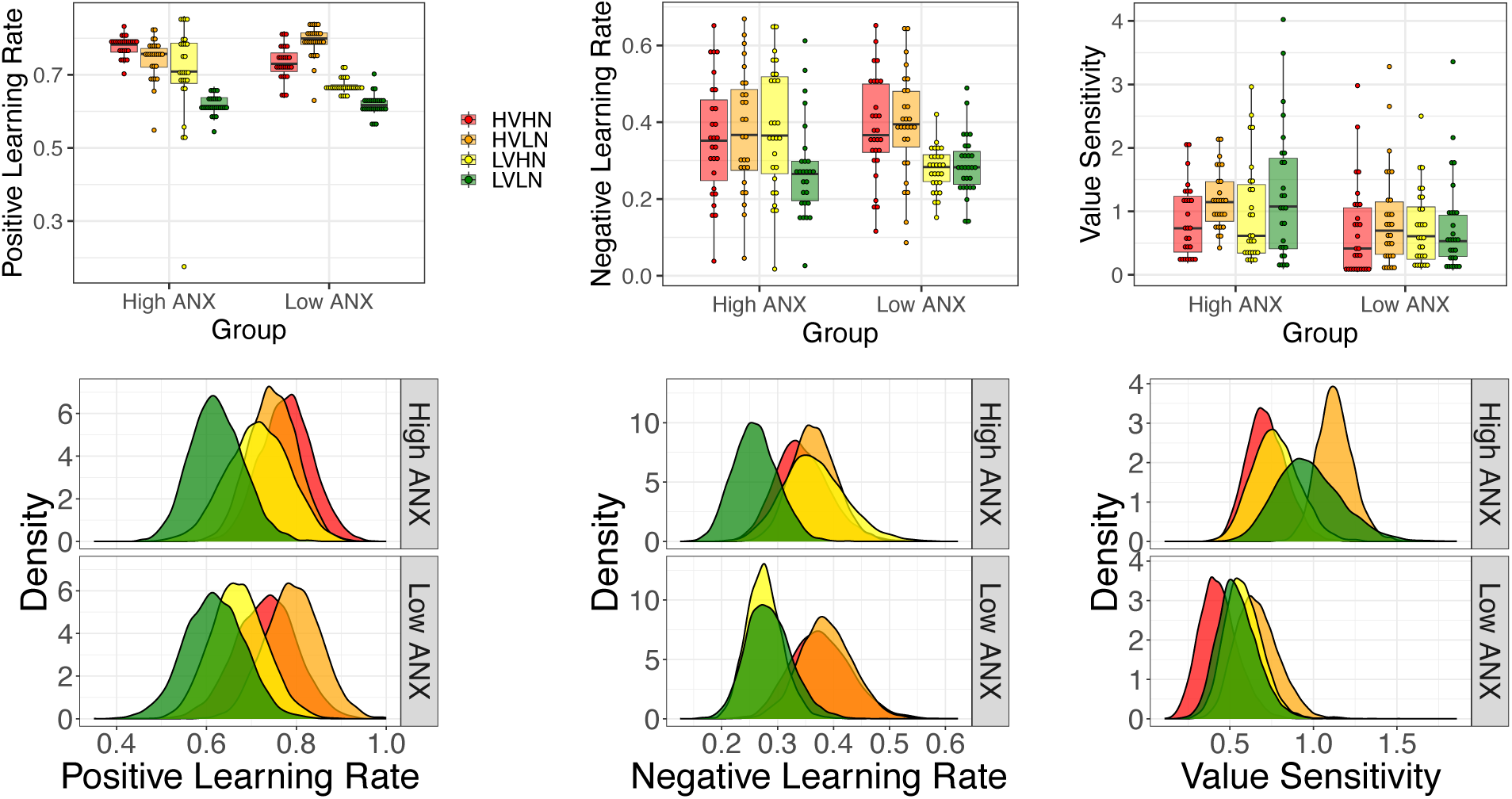
Alternative low and high ANX groups FU-RP models without indecision point. Individual-level parameter estimates and group-level distributions for positive and negative learning rates and value sensitivity parameters.

**S3 Fig.**
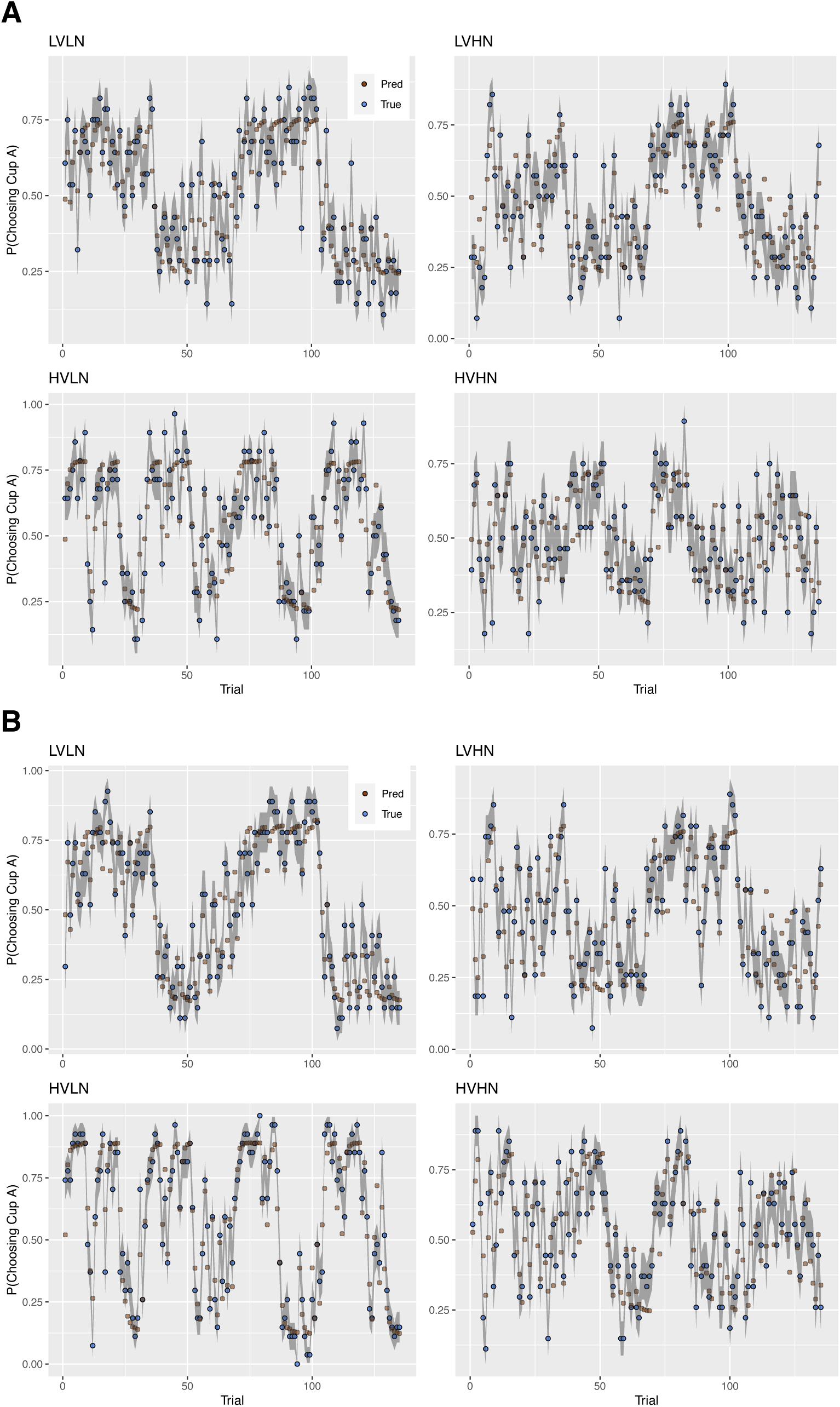
Model validation. Posterior predictive checks (PPCs) on reported hBayesDM models for **(A)** the low ANX group, and **(B)** the high ANX group. Predicted choices were found to largely track true choices across time, for each condition in each ANX group. Error bars represent 95% CIs around the mean.

**S4 Fig.**
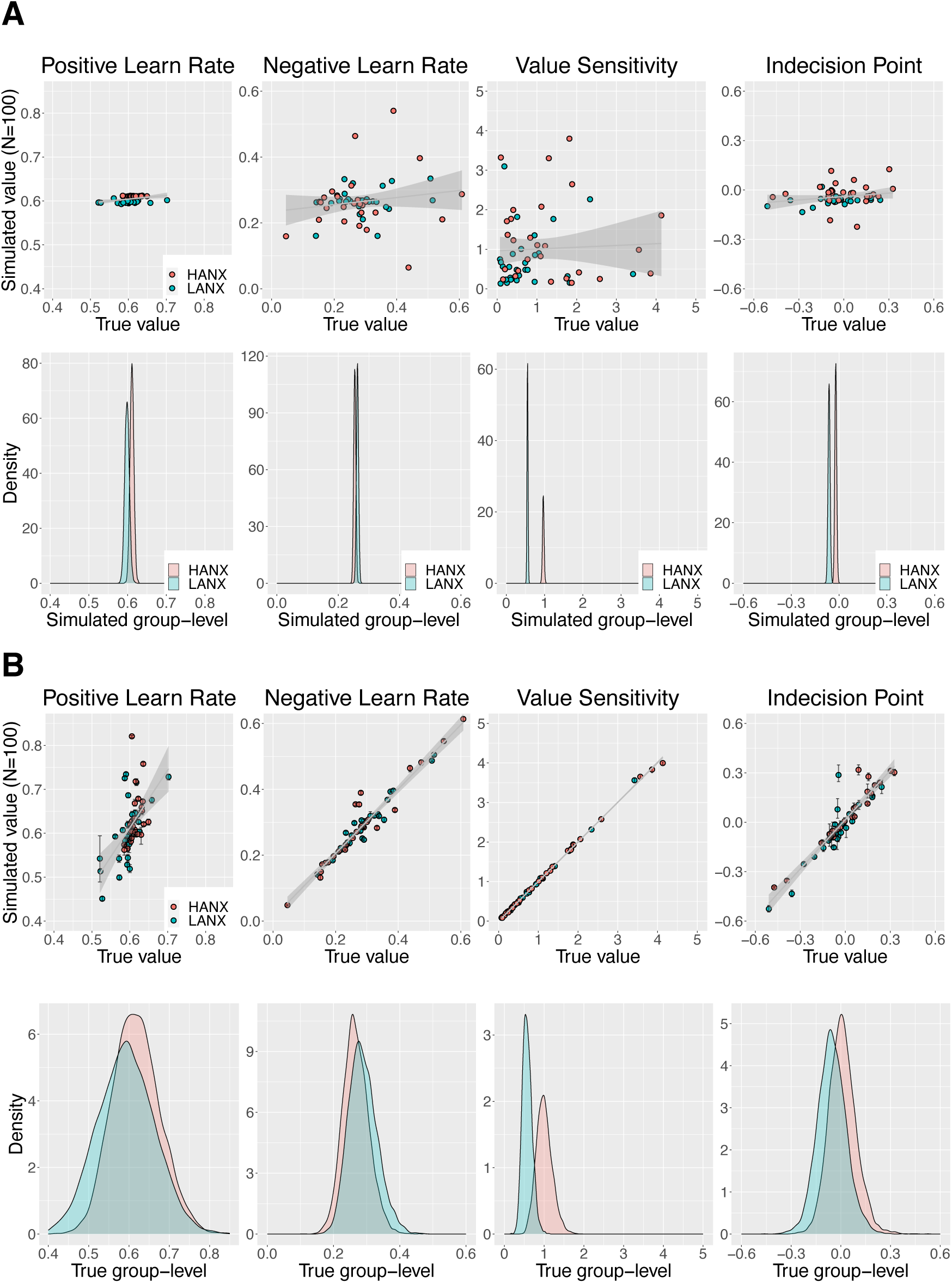
Parameter recovery. **A)** Simulation method 1: 100 synthetic datasets were generated per original participant and included in one large hierarchical model per low and high ANX group. Individual-level estimates were not recoverable, i.e., true and simulated values were not strongly correlated (upper panel). Group-level distributions were recoverable, however, and highly precise (lower panel). **B)** Simulation method 2: 100 simulated datasets were fit in a single hierarchical model per original participant. True and simulated individual-level parameters were highly correlated (r>0.9), except for a moderately-correlated positive learning rate (r=0.582) (upper panel). The true group-level posteriors (lower panel) show a relatively wider distribution for positive learning rate, likely constraining more variable individual-level estimates. Note that the same x-axis range is used for all plots of the same parameter for comparison purposes. Individual-level errorbars represent ±1 SD around the mean of the simulated estimates.

**S5 Fig.**
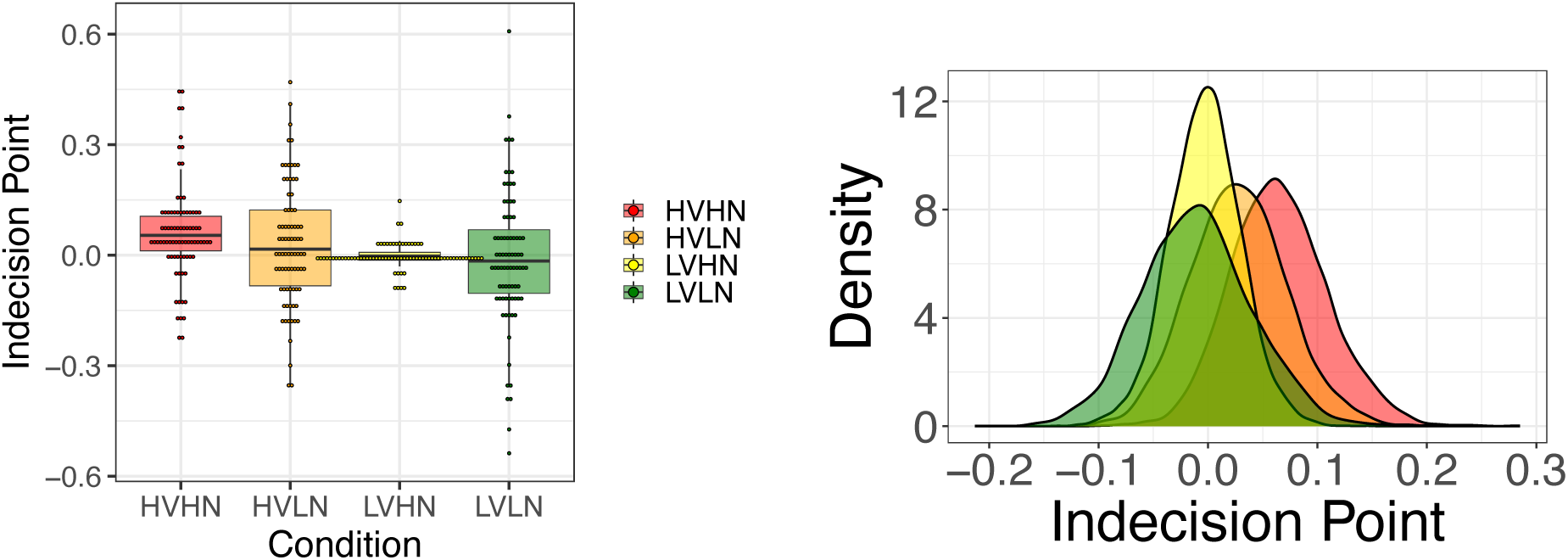
Reported FU-RP-IP model on full sample. Individual-level parameter estimates (left) and group-level distributions (right) are shown for the indecision point parameter.

**S6 Fig.**
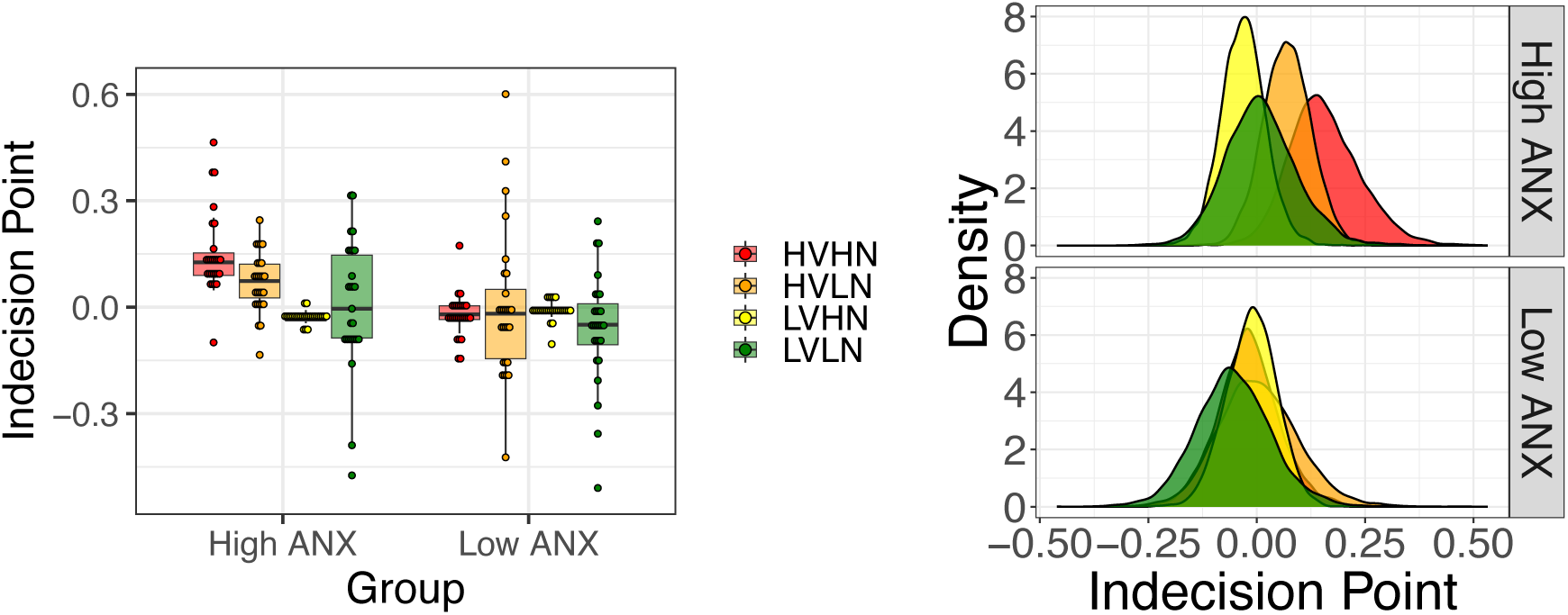
Reported FU-RP-IP model on low and high ANX groups. Individual-level parameter estimates (left) and group-level distributions (right) are shown for the indecision point parameter.

**S7 Fig.**
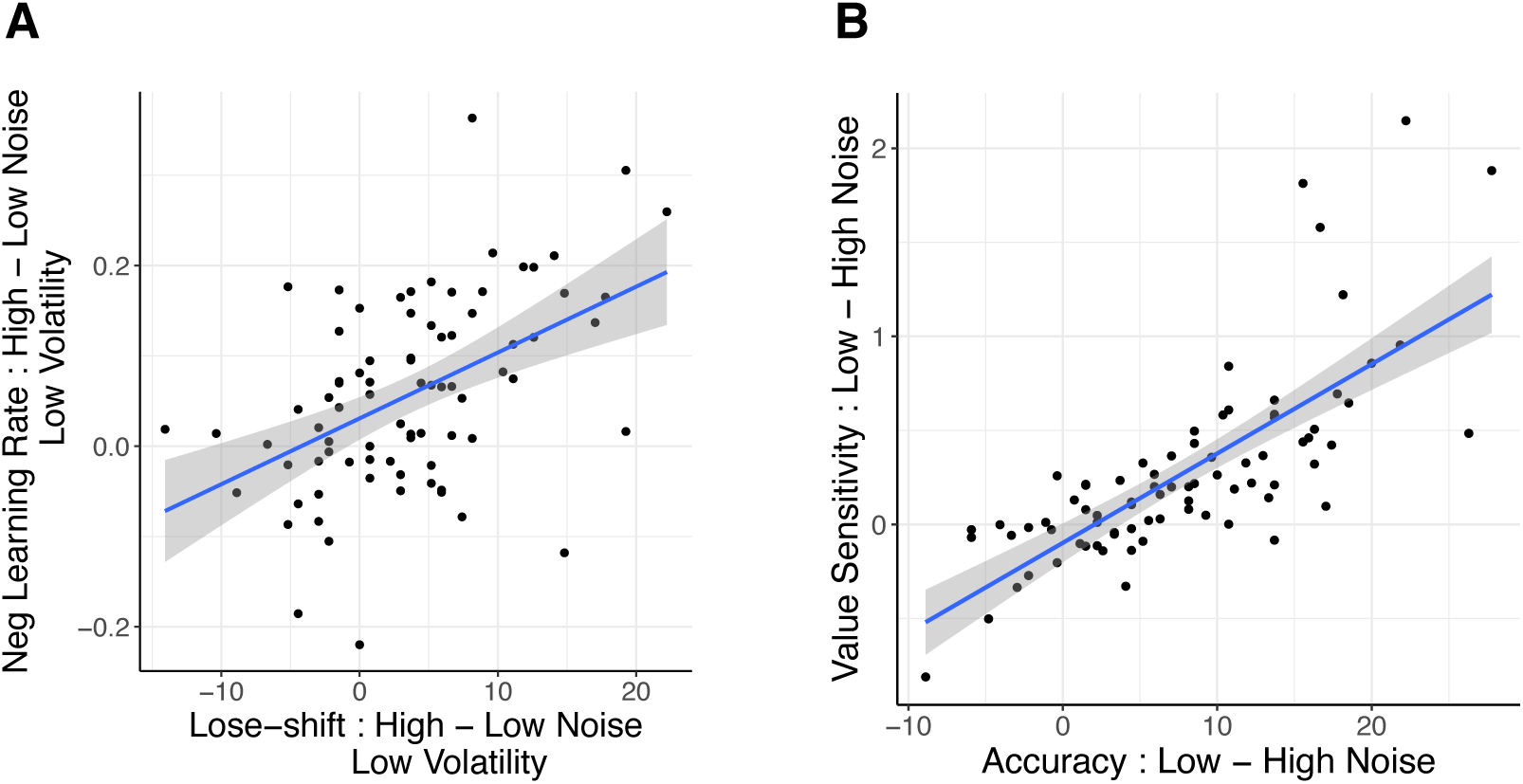
Relationship between behaviour and model learning parameters in full sample. A) Under low volatility, an increase in lose-shift behaviour from low to high noise conditions correlates with an increase in negative learning rate for the same comparison (r=.487, p<.001), and **B)** an increase in task accuracy from low to high noise conditions is associated with a similar increase in value sensitivity (r=0.751, p<.001)

#### Supplemental Tables

**S1 Table.**
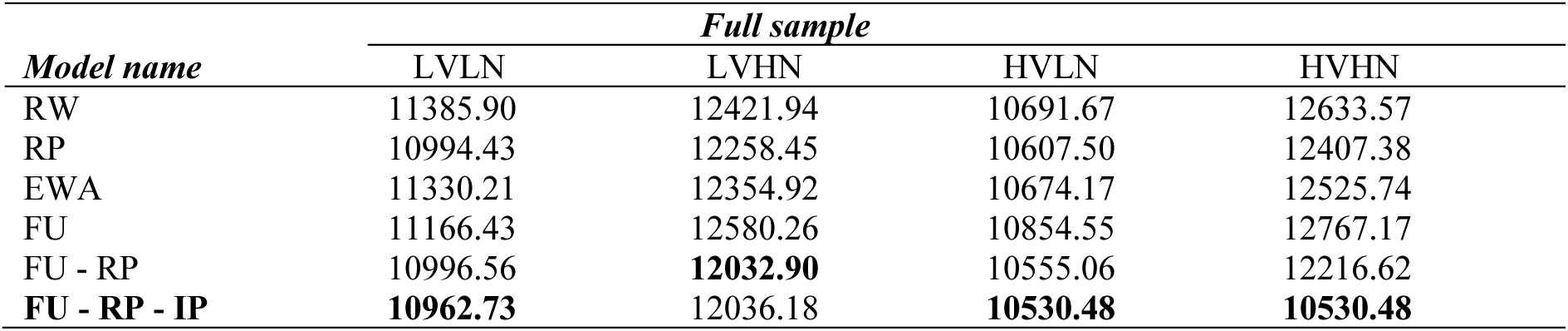
Model fits of full sample: LOOIC values. The best fitting model was the fictitious update model with both reward and punishment learning rates and indecision point parameters (FU-RP-IP). In fictitious update models, the expected value of the non-chosen stimulus also gets updated on each trial, based on the feedback received for the chosen stimulus. A similar model without the indecision point (FU-IP) was the best fit for the LVHN condition, although LOOIC values are close between the two models.

| *Model name* | *Full sample* |  |  |  |
| --- | --- | --- | --- | --- |
|  | LVLN | LVHN | HVLN | HVHN |
| RW | 11385.90 | 12421.94 | 10691.67 | 12633.57 |
| RP | 10994.43 | 12258.45 | 10607.50 | 12407.38 |
| EWA | 11330.21 | 12354.92 | 10674.17 | 12525.74 |
| FU | 11166.43 | 12580.26 | 10854.55 | 12767.17 |
| FU - RP | 10996.56 | **12032.90** | 10555.06 | 12216.62 |
| **FU - RP - IP** | **10962.73** | 12036.18 | **10530.48** | **10530.48** |

**S2 Table.**
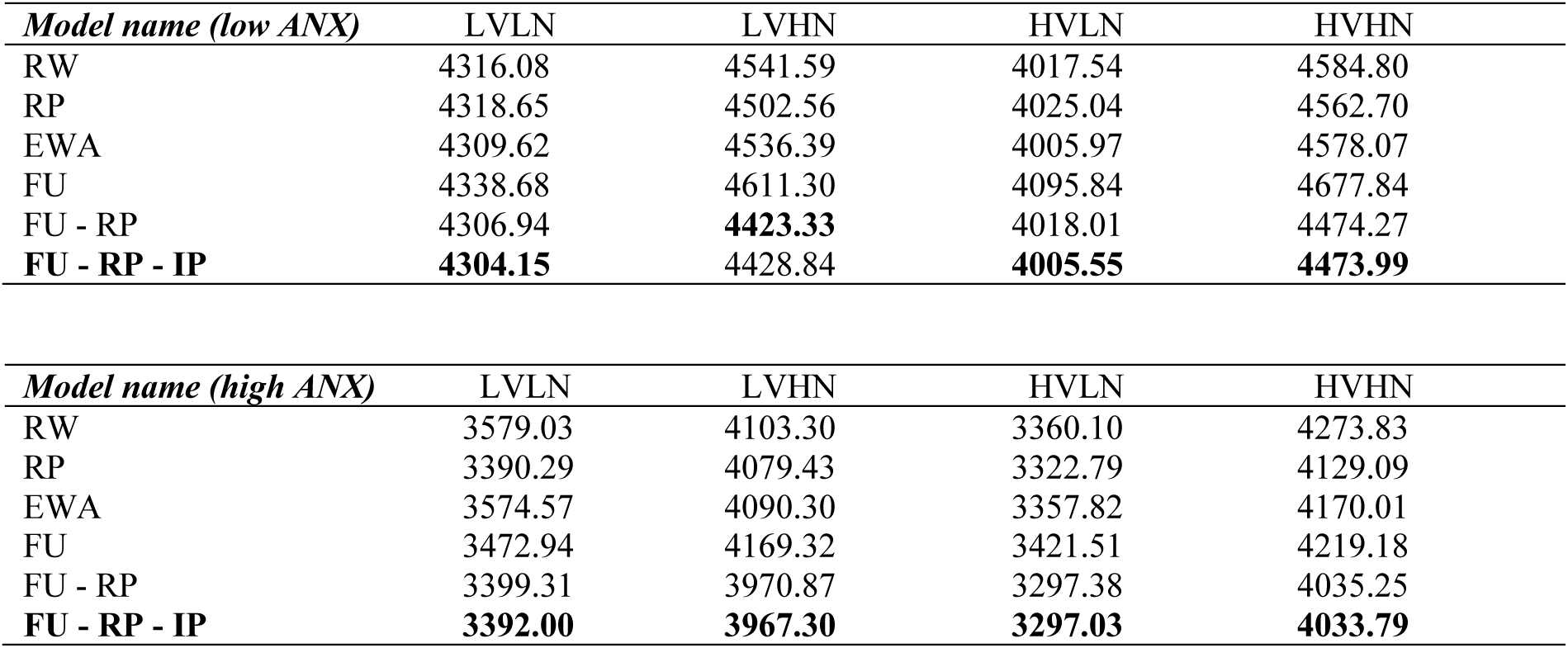
Model fits of low and high ANX groups: LOOIC values. The best fitting model was FU-RP-IP. The FU-IP model was best for the LVHN condition in the low ANX group only, although LOOIC values were only slightly higher for the FU-RP-IP model.

**S3 Table.**
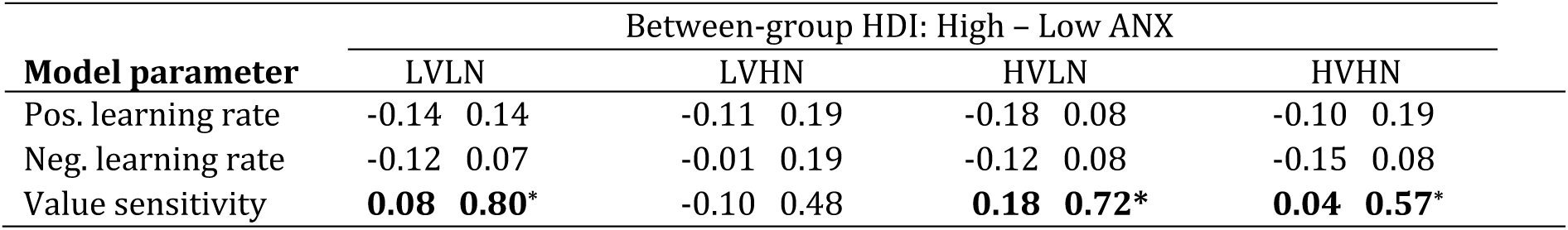
Alternative low and high ANX groups FU-RP models without indecision point. Differences in model parameter distributions between low and high ANX groups.

**S4 Table.**
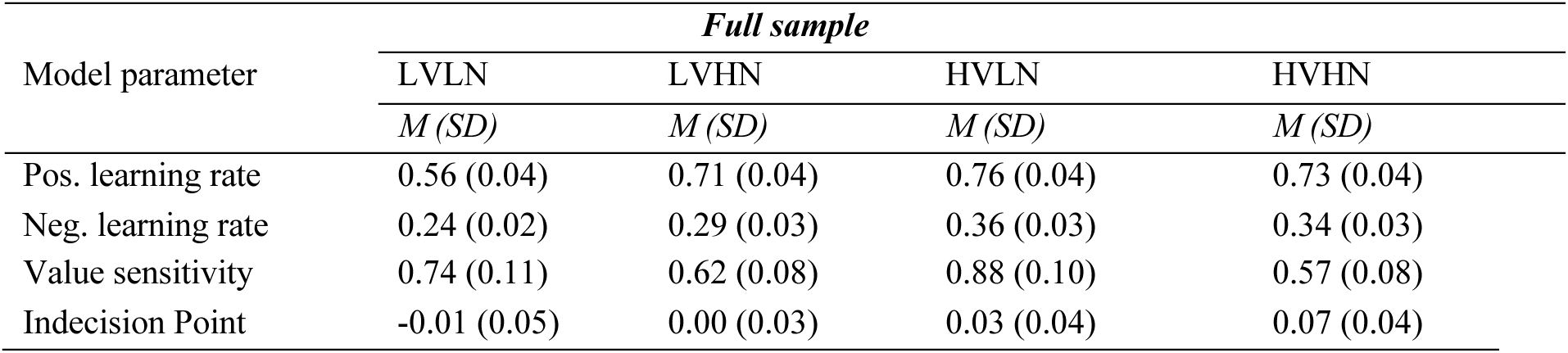
Median group-level parameters of reported FU-RP-IP model for the full sample.

**S5 Table.**
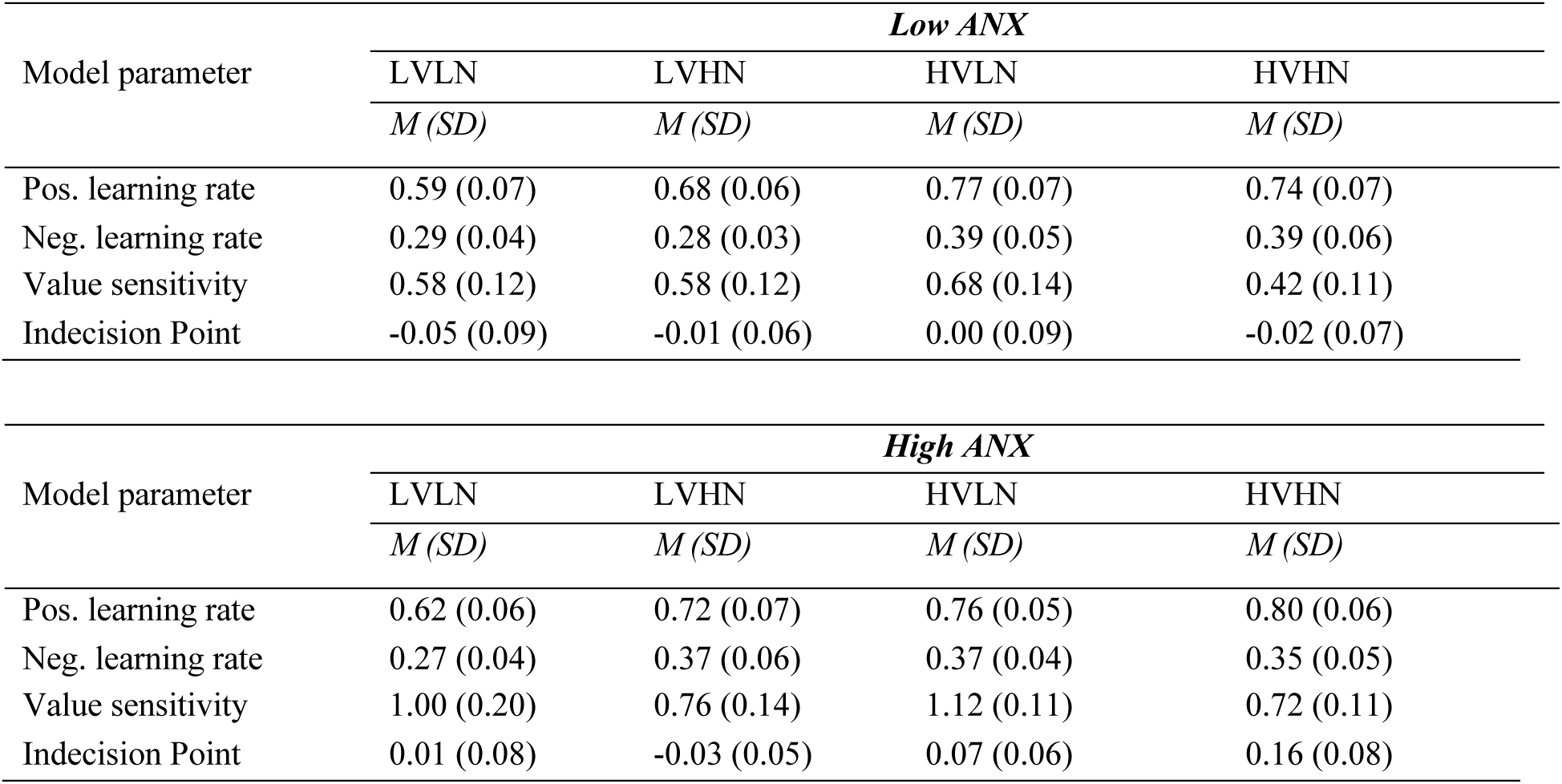
Median group-level parameters of reported FU-RP-IP model for low and high ANX groups.

## Notes

### Competing Interest Statement

The authors have declared no competing interest.

